# Iterative co-creation of harmonized human and non-human primate cellular and structural ontologies and 3D common coordinate frameworks for the basal ganglia

**DOI:** 10.64898/2026.07.30.741796

**Authors:** Song-Lin Ding, Ashwin A. Bhandiwad, Burke Q. Rosen, Stephanie C. Seeman, Brian Long, Nelson J. Johansen, Ugur Bayindir, Benjamin Facer, Yuanyuan Fu, Yoel Halimi, Yujie Hou, Dingzi Hu, Mike Huang, Takuro Ikeda, Phil Lesnar, Xiao-Ping Liu, Zhaoke Luo, Patrick L. Ray, Joshua J. Royall, Matthew T. Schmitz, Akiko Uematsu, Julien Vezoli, Faraz Yazdani, Trygve E. Bakken, Winrich A. Freiwald, Rebecca Hodge, Brian E. Kalmbach, Henry Kennedy, Lauren Kruse, Tyler Mollenkopf, Lydia Ng, David Osumi-Sutherland, Carol L. Thompson, Takuya Hayashi, Michael J. Hawrylycz, Matthew F. Glasser, David C. Van Essen, Hongkui Zeng, Ed S. Lein

**Author notes:** **Lead contact**: Ed S. Lein. **Corresponding authors**: Ed S. Lein and Song-Lin Ding.

## Abstract

A major goal of the BRAIN Initiative Cell Atlas Network (BICAN) is to create a suite of foundational reference cell atlases and associated standards for human and non-human primate brains. Central to this goal is the creation of cross-species harmonized cellular taxonomies and structural parcellations with formal ontologies that can be mapped into 3D reference frameworks bridging neuroimaging and cellular and histological resolutions. We describe here an iterative approach, focused initially on the basal ganglia, to co-create structural and cellular ontologies in human, macaque and marmoset brains, including a Harmonized Ontology of Mammalian Brain Anatomy (HOMBA), and to map and refine structural parcellations into neuroimaging-based common coordinate frameworks. These references provide the framework for documenting and mapping all experimental sampling in BICAN, allowing analyses of cellular and molecular variation as a function of topographic position, and enabling comparisons of cellular, molecular and neuroimaging-based functional variation within and between primate species.

**Highlights:**

- A hierarchical Harmonized Ontology of Mammalian Brain Anatomy (HOMBA) covering 2348 structures
- HOMBA-annotated 3D common coordinate frameworks (CCFs) of the basal ganglia across species
- Histologically informed 3D parcellation/atlas of 280 human subcortical structures indexed by HOMBA
- Mapping and integration of structural, cellular and functional data with HOMBA and CCFs

## INTRODUCTION

Single cell and spatial genomics methods are rapidly transforming neuroscience and many other fields of biology and medicine. The NIH BRAIN Initiative Cell Census project (BICCN) has spearheaded consortium-scale efforts to establish robust single cell genomics methodologies and to apply them across the entire brain to produce whole brain cell atlases in mouse (Langlieb et al. 2023; Yao et al. 2023; Zhang et al. 2023) and a draft cell atlas of the human brain (Siletti et al. 2023) in the BICCN phase (RRID:SCR_015820). The current phase of this effort, the BRAIN Initiative Cell Atlas Network (BICAN, RRID: SCR_022794), is designed to move from a draft to a comprehensive cell atlas in human and non-human primate (NHP) brains, and to produce a harmonized set of foundational references in the spirit of the human genome project.

The Human and Mammalian Brain Atlas (HMBA) project aims to create the foundational adult brain atlases for BICAN. These atlases contain a series of core components, all designed to allow harmonization of homologous elements across human, macaque and marmoset brains (and eventually mouse and other mammalian species): 1) single cell genomics-based cell classifications and formal cell ontologies and nomenclatures, 2) anatomical parcellations and ontologies informed by cellular spatial organization, and 3) 3D structural parcellations on neuroimaging-based probabilistic templates, or common coordinate frameworks (CCFs). Remarkable work in the mouse brain serves as a template for this work, albeit a relatively simple one given the size and stereotypy of the adult mouse brain. Whole mouse brain single cell genomics-based cell taxonomies are complete, and they define over 5000 types of neurons and non-neuronal cell types (Yao et al. 2023). Whole mouse brain spatial transcriptomics (STx) has been performed to map the spatial distributions of those cell types (Yao et al. 2023), and those spatial distributions have been mapped into a probabilistic 3D atlas, the Allen Mouse Brain CCFv3 (Wang et al. 2020).

Creating these core components in human and non-human primate brains, and harmonizing across species, is much more challenging given the larger size and greater interindividual and interareal variability of primate brains. Multiple 3D reference atlases currently exist in each species, but most NHP atlases are based on single brains, and all have different structural delineations and nomenclatures (Hartig et al. 2021; Saleem et al. 2021; Saleem et al. 2024). Furthermore, the time constraints of the BICAN project necessitated the concurrent co-creation of different elements, followed by an iterative strategy for refinement. Available tools and resources made this feasible: 1) A whole brain adult human histologically informed reference (Ding et al. 2016) with a brain-wide hierarchical cell taxonomy (Siletti et al. 2023), plus a 3D parcellation on the MNI152 template (Ding, Royall, 2020) that provided a starting point for the entire effort; 2) A draft whole human brain single-cell genomics-based cell atlas (Siletti et al., 2023), and multiple comparative studies in human, macaque, marmoset, and mouse (Hodge et al., 2019; Bakken, Jorstad, et al. 2021; BICCN, 2021) demonstrating the feasibility of creating detailed cell taxonomies in specific brain regions and mapping homologous cell types across species; 3) Successful use of spatial transcriptomics methods to map the spatial distributions of cell types in the most challenging human brain tissues (Jorstad et al. 2023; Gabitto et al. 2024); 4) Ongoing efforts to create the next generation of probabilistic MRI-based templates in human, macaque, and marmoset brains that could serve as the core CCFs for BICAN as a whole (Hayashi et al. 2021).

These tools allowed a pragmatic, iterative co-creation strategy for cross-species harmonized cell atlas creation, initially focused primarily on the basal ganglia as a well-defined region to establish a paradigm that can be extended to the entire brain. These reference frameworks interact with many of the other studies describing parts of the overall BICAN effort. The adult human brain atlas and structural ontology (Ding et al. 2016) was used to create a Harmonized Ontology of Mammalian Brain Anatomy (HOMBA), which serves as the common language for anatomical structures for groups in BICAN to analyze with single cell or spatial genomics methods, and as an ontological element for sample documentation and tracking across BICAN. The structural nomenclature in HOMBA was then used as a key element in the cell type nomenclature system to describe anatomical location/distribution. Structural parcellation based on HOMBA was then used to annotate new probabilistic templates, or CCFs, across species in 3D. Mapping spatial transcriptomics data into the CCFs closed the loop to allow comparisons and refinement of HOMBA structural boundaries and create the final cell atlases: CCFs in each species, parcellated with a cellularly-informed harmonized structural ontology allowing the mapping of cellular spatial distributions with harmonized cell ontologies and nomenclatures. This approach sets the paradigm for the rest of the brain, with the first steps of creating HOMBA and 3D human structural parcellations for the rest of the brain outside of the neocortex well under way.

## RESULTS

### Overall workflow for creating harmonized structural ontology and 3D CCFs

The overall goals of HMBA are to create a harmonized structural and cellular atlas of the human, macaque, and marmoset brain. This atlas consists of 3 major data components (**Fig. 1**): 1) A comprehensive classification of cell types derived from 10x Multiome (combined RNA-seq and ATAC-seq) profiling, generated in parallel and integrated across species (Johansen et al. 2025) (**Fig.1A**), 2) Spatial tissue maps of cell types with spatial transcriptomics methods to map topographic distributions and local tissue microarchitecture (Hewitt et al., 2025) (**Fig.1B**), and 3) mapping and integration of multimodal data with harmonized HOMBA ontology and CCFs (**Fig. 1C**). Structural and cellular ontologies and nomenclatures provide essential linkages across these components and allow common cross-species structural parcellation and spatial mapping of cellular distributions in 3D space to bridge scales.

**Figure 1.**
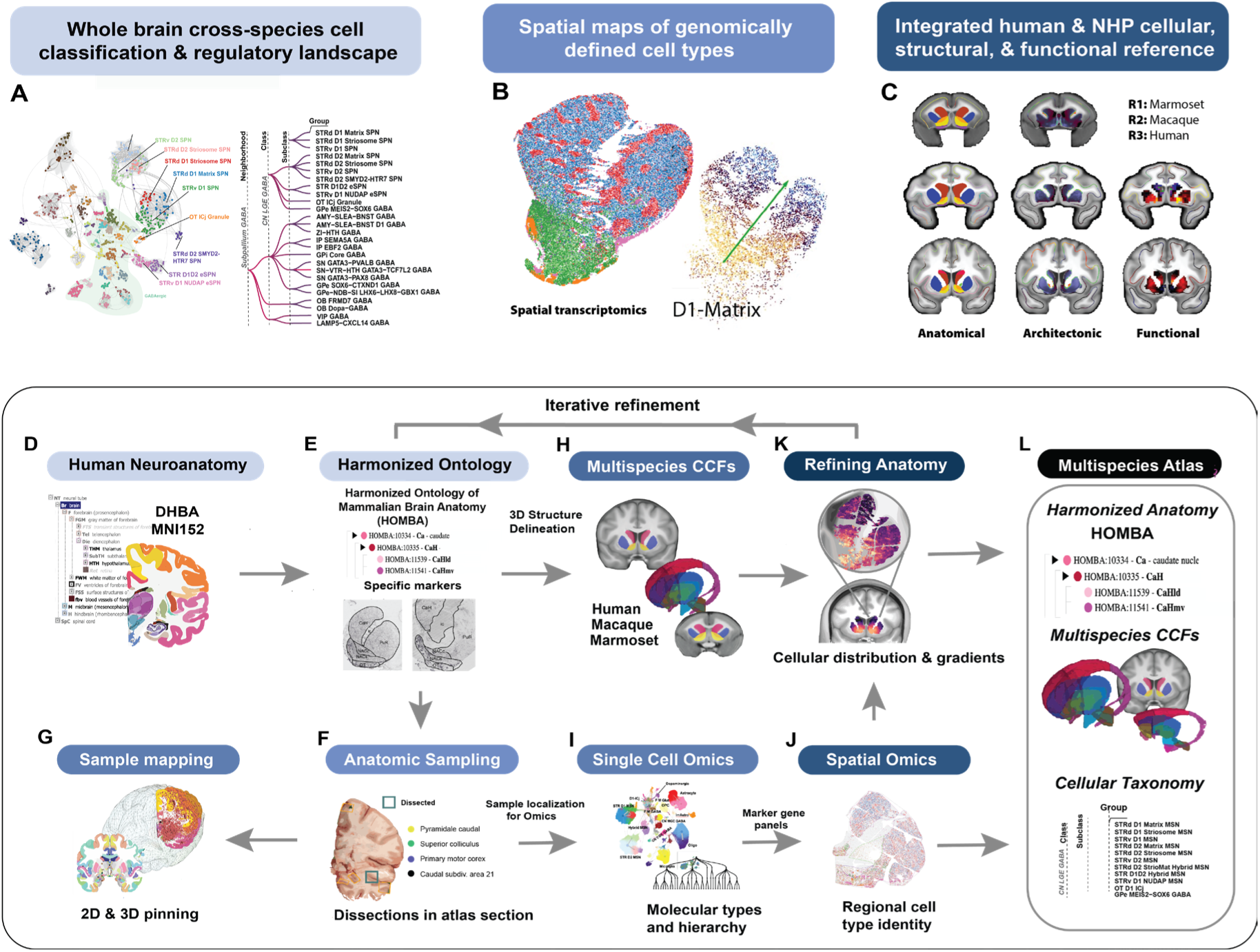
Major components and workflow. (**A**) Whole brain cross-species cell type classification, with cell type UMAP embedding and hierarchical organization. (**B**) spatial maps of genomically defined cell types, with PCA of expression indicating spatial gradient in D1-Matrix. (**C**) Integrated human and NHP anatomical, architectonic, and functional images. Marmoset, Rhesus Macaque, and Human coronal volume slices illustrating the BG. Left column: HMBA BG structural parcellation. Middle column: T1w/T2w and cortical myelin map. Third column: fMRI ICA components, individual in Macaque, group in Human (HCP1071) and omitted in marmoset. (**D-L**) Iterative co-creation of cross-species anatomical CCFs with single cell and spatial transcriptomics-based characterization of anatomically anchored cell types (see text for detailed workflow).

The atlas creation workflow is presented in **Fig.1D-L**. The structural HOMBA ontology was a critical early element in creation of the atlas, necessary not only to describe anatomical divisions but to drive coordinated cross-species sampling for single cell analysis. The Developing Human Brain Atlas (DHBA) ontology (**Fig.1D**) served as a starting point, based on whole brain adult and developmental 2D and 3D atlases (Ding et al. 2016, 2022; Ding, Royall, et al. 2020). DHBA was extended to create HOMBA, harmonizing structural nomenclatures and structural parcellations across human, macaque and marmoset (**Fig.1E**). These parcellations were used to both drive physical dissections for snMultiome analyses (**Fig. 1F**), and to pin sample locations onto 2D atlas plates and 3D CCFs (**Fig. 1G**). The HOMBA ontology was used to label 3D parcellations of brain structures in human, macaque and marmoset CCFs to create harmonized 3D atlases for these species (**Fig. 1H**). snMulitome profiling was used to create the cross-species harmonized cell ontology (Johansen et al. 2025) (**Fig. 1I**), and spatial transcriptomics data was generated in each species to map the cellular distributions in sections spanning all structures (Hewitt et al., 2025) (**Fig. 1J**). Spatial transcriptomics sections were then mapped into the 3D CCFs and used to refine boundaries based on cellular distribution patterns where justified (**Fig. 1K**). The final product, described here for basal ganglia structures, is a cross-species harmonized structural ontology (HOMBA), 3D atlases consisting of HOMBA-annotated CCFs, and a cellular ontology whose nomenclature incorporates the HOMBA structures where those cells are located (**Fig. 1L**).

It should be mentioned that the basal nuclei (BN) typically consists of striatum (STR) and globus pallidus (GP: GPe plus GPi), whereas the basal ganglia (BG) typically also includes the subthalamic nucleus (STH) of the diencephalon, and substantia nigra (SN) and ventral tegmental area (VTA) of the midbrain. In this study, we have included the STH, SN, and VTA as connectionally and functionally related structures as part of BG. Thus, all of these components (STR, GP, STH, SN, VTA) are grouped as a BG system (Joseph and Cardozo, 2004) for the current study, rather than a hierarchical term in the HOMBA ontology.

### Generation of a 3D human brain atlas for sample pinning and documentation

To establish the approach for creating 3D structurally annotated MRI-based templates that could be used to facilitate accurate pinning and documentation of whole human brain tissue samples used for single nucleus genomics profiling in 3D space, we first reconstructed a 3D whole human brain atlas (**Fig. 2A-D**) on the ICBM2009b MNI152 average template (the MNI152 template for short) based on our 2D histological human brain atlas and associated structural ontology [i.e., DHBA ontology; see (S.-L. Ding et al. 2016, 2022); **Fig. 2A,B**] as well as *in situ* hybridization (ISH) data from the Allen Human Atlas (Hawrylycz et al. 2012). For example, three closely adjacent human thalamic sections for the gene markers *CHRM3*, *CARTPT* and *CBLN2* can be aligned to enable accurate 2D delineation of the thalamic nuclei (**Fig. S1**) and then mapped to 3D template space. In the 3D MNI152 template, all cortical gyri and major subcortical structures including BG were 3D-reconstructed based on 2D histological delineation (**Fig. 2C, D**; see (Ding, Royall, et al. 2020).

**Figure 2.**
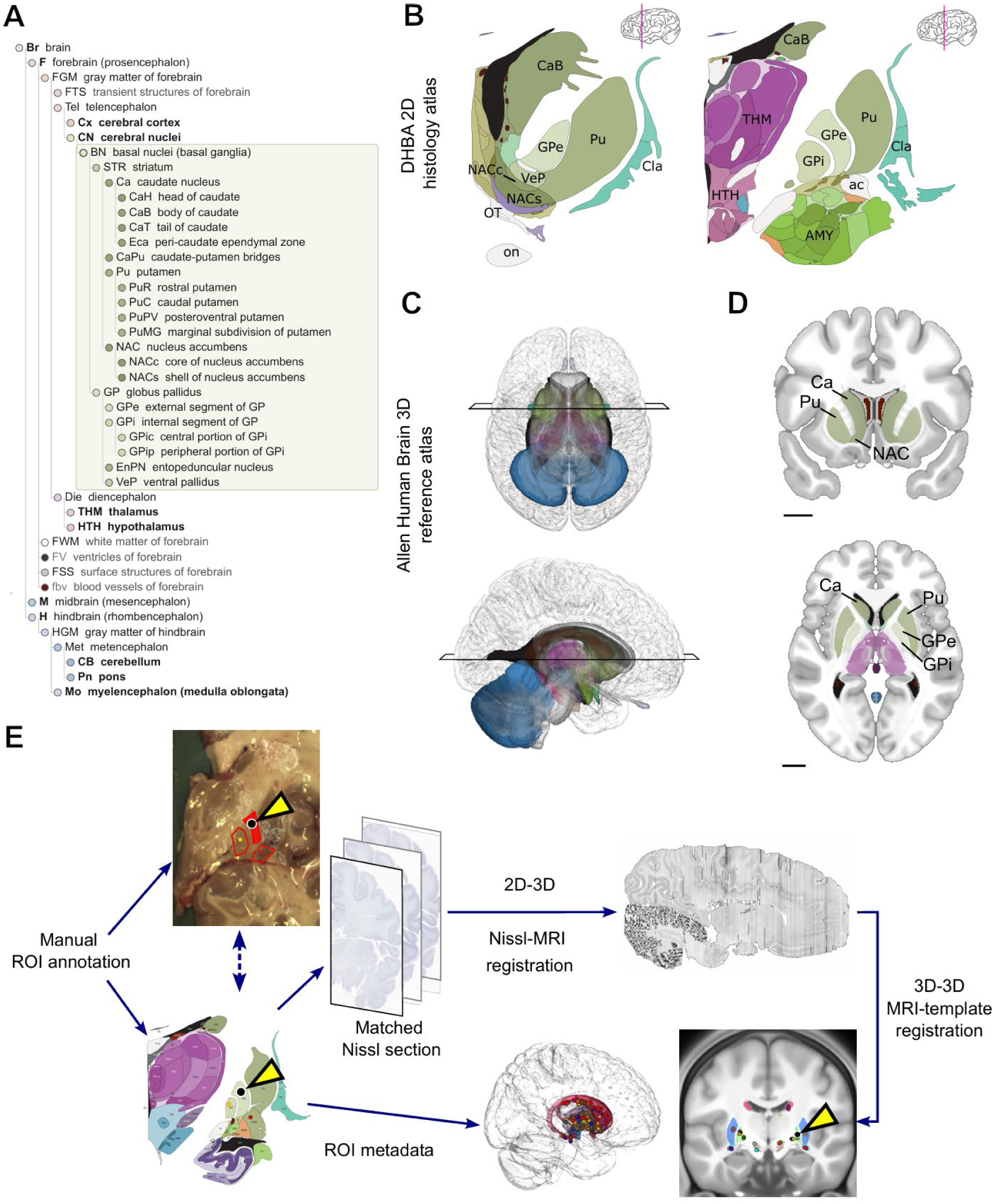
Single cell RNA-seq sample pinning onto 3D human brain atlas. (**A**). Organization of the DHBA structural ontology with basal nuclei (basal ganglia) expansion. Structures are divided into forebrain, midbrain, and hindbrain divisions and further divided into gray matter, white matter, ventricles, surface structures, and blood vessels. Bolded labels show major divisions of the brain. (**B**). Example sections of the 2D DHBA plate atlas. (**C,D**). 3D and 2D representations of subcortical structures in the Allen Human Brain 3D reference atlas on the MNI152 template with BN (BG) structures labeled. (**E**). Schematic diagram describing sample pinning protocol, based on (Casamitjana et al. 2022). Sampling ROIs are pinned manually in the slab image and independently in the matched section of the DHBA 2D plate atlas (arrowheads). Dashed line signifies correspondence between coordinate spaces. 2D plate atlas sections are registered to a 3D MRI coordinate space using the matched Nissl template section. A final registration places the pins in the HCP MNI coordinate space. An interactive 3D visualization can be found at https://allen.neuroglass.io/glances/06a20bab-c01e-7cf9-8000-6c034a3597fb

As reported previously, 2D delineation of brain structures on histological sections can be manually mapped to 3D MRI space (Scholtens et al. 2018; Pijnenburg et al. 2021). Once consistent and accurate delineations of target brain structures are established in 2D sections (e.g., **Fig. S2**), these delineations can be mapped and/or projected to different 3D MRI templates to generate 3D atlases using ITK-SNAP (Yushkevich et al. 2006; Wang et al. 2020). In the present study, we delineated the 3D MNI152 template in two stages. First, structures that are easily identified on the MNI152 template were directly parcellated using ITK-SNAP. In this case, most of the large subcortical structures (e.g., basal ganglia, amygdala, thalamus, hypothalamus, cerebellum, pons and medulla) are clearly identifiable based on intrinsic signals in the MRI template (e.g., contrast and intensity) and anatomical landmarks (major sulci and other landmarks; see (Petrides et al. 2012; S.-L. Ding et al. 2016; ten Donkelaar et al. 2018).

A secondary process was used to delineate structures or subregions which are not clearly distinguishable in the template. This method involved matching histological sections to high- and/or low-resolution MRI slices with guidance by landmarks, anatomical topography, and intrinsic signals in the template. For example, based on the locations and topography of the major amygdaloid nuclei (e.g., basolateral nucleus: BL, basomedial nucleus: BM, lateral nucleus: La, and posterior cortical nucleus-periamygdaloid cortex: CoP-PAC, in **Fig. S2A**’) in histological sections, we delineated these amygdaloid structures in the matched high-resolution 7T MRI images (T2w) from a single brain (Edlow et al. 2019; an amygdala slice is shown in **Fig. S2A**), which is registered to the low-resolution MNI152 template (T1w in **Fig. S2B**). Then we drew the boundaries and/or projected the drawings (with slight modification) to the matched MNI152 template slice (**Fig. S2B**). Within each major amygdaloid nucleus such as BL, its subdivisions were further segmented based on matched histological sections, which display defining features of the subdivisions such as overall cell sizes (BLmc containing mostly large cells, BLi mostly medium-sized cells and BLpc mostly small cells; **Fig. S2A**’) and/or subregion-specific gene expression (e.g., BLmc and BLi containing strong and moderate non-phosphorated neurofilament protein (NFP) expression, respectively, while BLpc has weak NFP expression). Similarly, 3D segmentation of human thalamic nuclei (**Figs. S2**D’, and **S2**D-F), major white matter bundles (**Figs. S2**G’ and **S2**G-I), and others was achieved in the MNI152 template. In total, 280 subcortical structures plus all cortical gyri were 3D parcellated on the MNI152 template (available via https://alleninstitute.github.io/CCF-MAP/descriptions/human_ccf.html).

Finally, the 2D DHBA atlas Nissl images (Ding et al., 2016) were mapped into the MNI152 template using a probabilistic registration model (Scholtens et al. 2018; Pijnenburg et al. 2021; Casamitjana et al. 2025) (**Fig.2E**). This transform provided a pragmatic way to map dissected samples into the 3D CCF. Slab images of brains being sampled for single nucleus genomics were compared to 2D reference atlas plates to identify the closest plane-matched 2D reference atlas plates, regions to be dissected were drawn on the slab images to document their location, and closest approximate locations were pinned to the reference atlas plate. The mapping of the underlying Nissl images to the MNI152 template described above provided the transform to map those sample locations in the MNI152 3D coordinate space. All sample dissections for single nucleus genomics analyses were drawn and documented in this fashion.

### Creating a harmonized ontology of mammalian brain anatomy (HOMBA)

The larger goal of HMBA is to create harmonized 3D annotated CCFs in human, macaque and marmoset. A key element of this goal was to create a common structural ontology and anatomical parcellation of homologous regions. We approached this by using a wide range of available informative histological, ISH and immunohistochemistry (IHC) data along with other available information across mammalian species (**Figs. 1E, S1, 3A-I, S3, S4**) to parcellate 2D histological images similarly in each species. Comparison of these spatial data across multiple species and sources permitted identification and boundary definition of homologous regions and boundaries. While these technically remain putatively homologous structures, the high degree of similarity and commonality of structural organization, gene expression patterns and literature across species (mirrored in cellular distributions as below) strongly support their evolutionary and functional homology.

**Figure 3.**
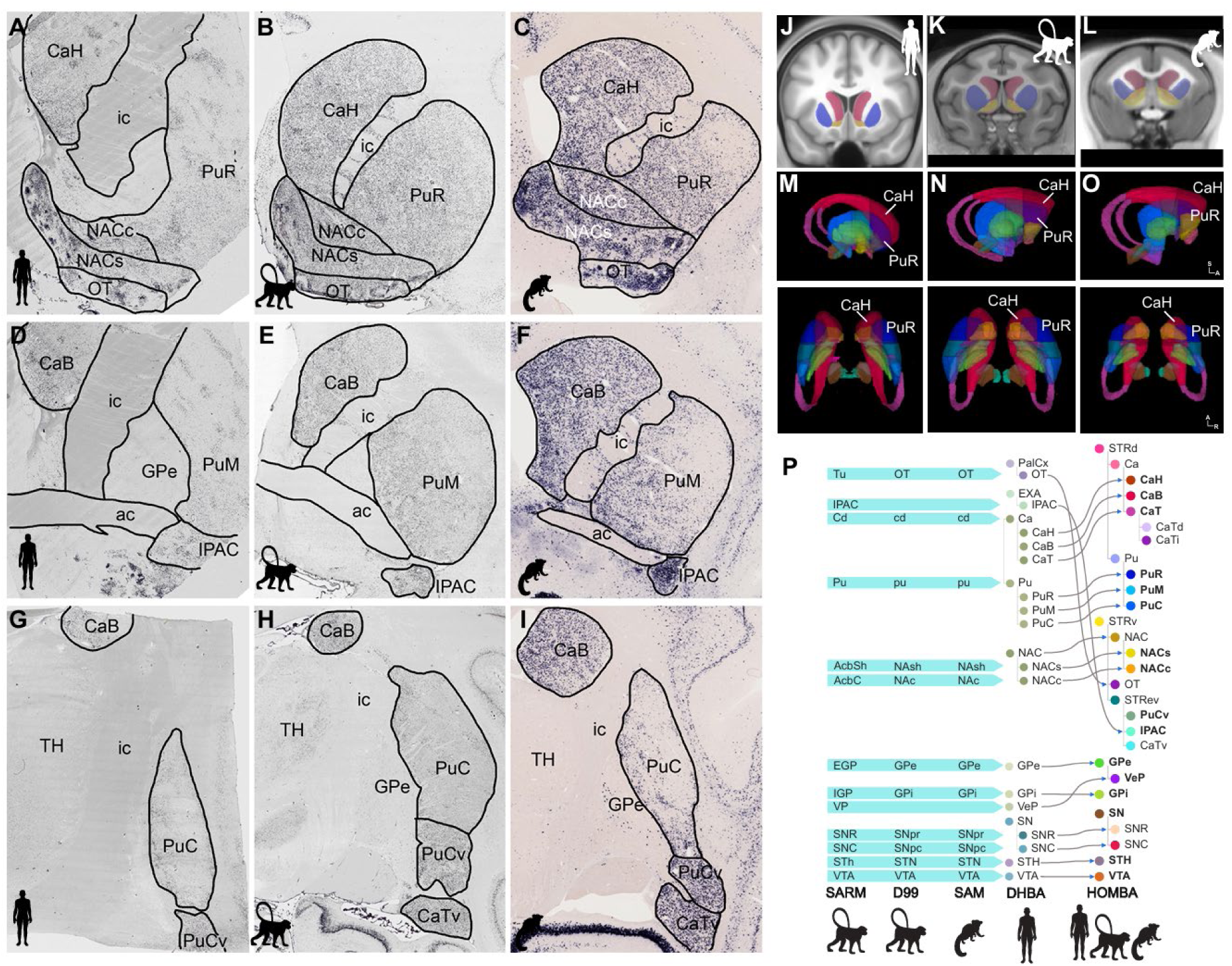
Creation of a harmonized ontology of mammalian brain anatomy and basal ganglia parcellation. **(A-I)** Examples of prodynorphin (*PDYN*) expression used for histological parcellation of BN/BG regions in human, macaque, and marmoset along the anterior (A-C) to posterior (G-I) axes. *PDYN* expression enables consistent demarcation of caudate and putamen subregions across species. (**J-O)** Equivalent brain region parcellations were delineated in species-specific three-dimensional spaces on reference templates shown in coronal T1w MRI slices (J-L) and in views from the side (top row) and from above (bottom row) of segmented and colored BG parcels (**M – O**). (**P)** The harmonized ontology of mammalian brain anatomy (HOMBA) was created based upon an initial framework of the Allen Human Reference Atlas (3D, v1.0; see Ding, Royall et al., 2020) with the DHBA ontology, and nomenclature was harmonized to existing reference atlases including the Subcortical Atlas of Rhesus Macaque (SARM), the D99 rhesus macaque atlas, and the Subcortical Atlas of Marmoset (SAM).

The striatum (STR) is a large and complex structure with great diversity in anatomy, gene expression and function (Haber 2016; Graybiel and Matsushima 2023), consisting of dorsal (STRd) and ventral (STRv) parts with STRd containing caudate (Ca) and putamen (Pu), and STRv containing nucleus accumbens (NAC) and olfactory tubercle (OT). To map these structures we combined and analyzed ISH and IHC data (e.g., **Fig. 3A-I**) as well as consistent anatomical landmarks (e.g., anterior commissure and caudal edge of the Pu across species). These data mapped gross structures, and also often illustrated gradient organization within structures as well. For example, complex medial (Ca)-lateral (Pu), ventral (STRv)-dorsal (STRd), and rostral-caudal differences/gradients in gene expression can be observed with specific gene markers across species (**Figs. 3, S3 and S4**). For instance, *PDYN* expression is overall strong and more homogeneous in Ca, but weaker and patchy-like in most parts of Pu (medial-lateral difference) except the rostral Pu, which shows stronger *PDYN* expression (**Fig. 3A-I**). This *PDYN* expression pattern is overall conserved in human (**Fig. 3A, D, G**), macaque (**Fig. 3B, E, H**), marmoset (**Fig. 3C, F, I**), and mouse (see Ma et al., 2025) brains. Other markers display obvious dorsal-ventral difference/gradients in STR [e.g., Wolfram syndrome 1 homolog (*WFS1)-ISH;* **Fig. S3**], rostral-caudal difference/gradient in Pu (e.g., calbindin-d28k (CB)-IHC; **Fig. S4**) or medial (weaker)-lateral (strong) differences in STR (e.g., *CNR1* expression*, data not shown*).

This analysis strategy (see examples in **Figs. S1, 3**A-I**, S3, S4**) enabled generation of a data-driven Harmonized Ontology of Mammalian Brain Anatomy (HOMBA) consisting of 2348 hierarchically organized anatomical terms spanning the entire brain and spinal cord (see HOMBA online link at https://alleninstitute.github.io/CCF-MAP/docs/HOMBA_ontology_v1.html). HOMBA was then used for labeling of 3D reconstructed BG components in all three species (**Fig. 3J-P**; see below **Fig. 4**).

**Figure 4.**
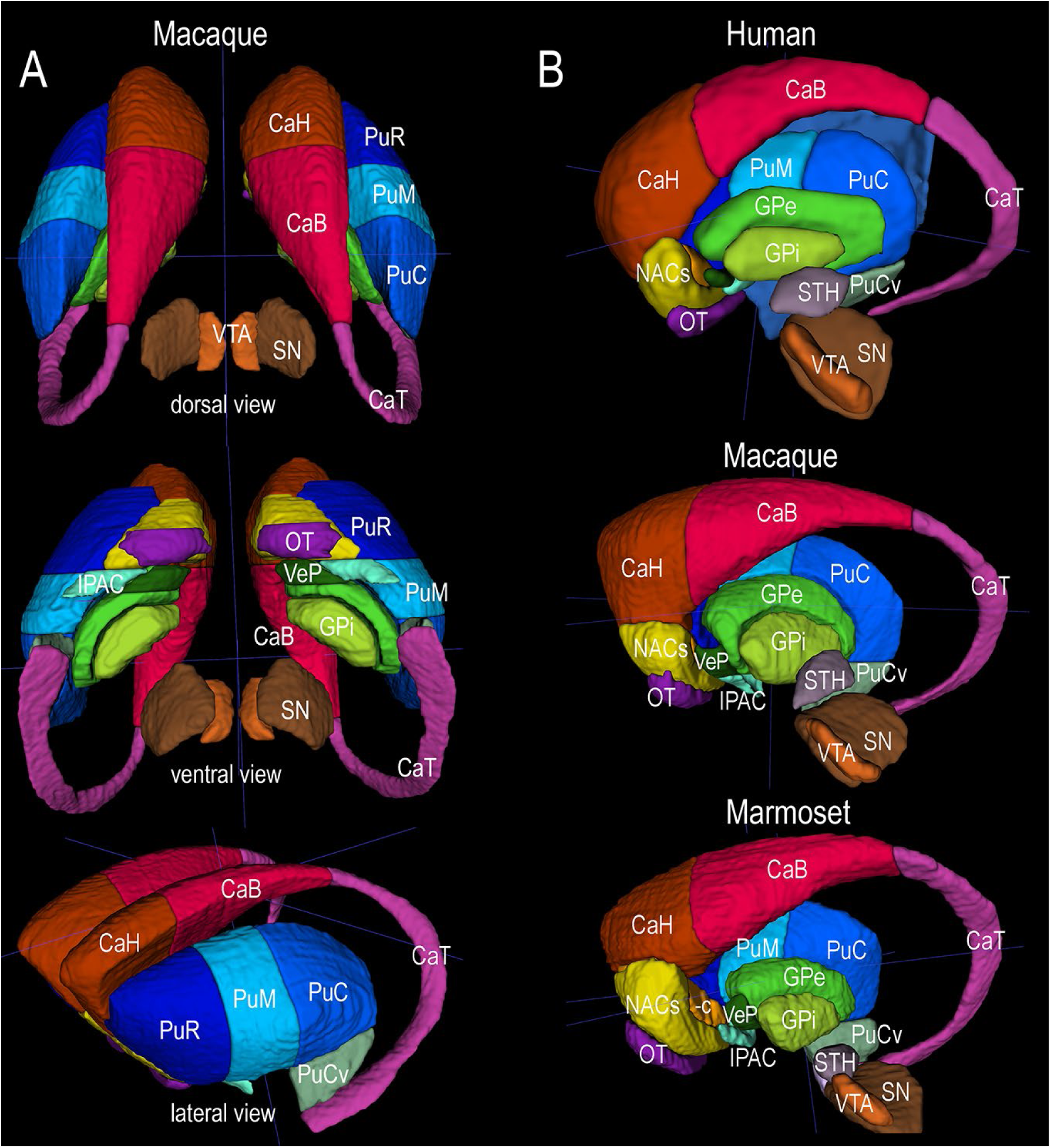
Harmonized 3D reconstruction of all BG subdivisions across species using HOMBA. (**A**) Dorsal, ventral and lateral views of 3D BG atlas/CCF of the macaque (from the top to bottom). The BG structures in both hemispheres are shown. (**B**) Medial views of 3D BG atlases/CCFs of the human, macaque and marmoset (from the top to bottom). In these harmonized BG atlases/CCFs across species, the same structures are coded with the same colors to facilitate the comparison. Detailed BG subdivisions reconstructed in 3D templates across species include three caudate (CaH, CaB and CaT), three putamen (PuR, PuM and PuC) and several ventral striatal (NACs, NACc, OT, IPAC and PuCv) subdivisions, as well as three subdivisions of globus pallidus (GPe, VeP and GPi). In addition, three closely related structures (STH, VTA and SN) are also 3D reconstructed.

Terminology in the HOMBA ontology was mainly derived from the DHBA human brain ontology described above and harmonized with other ontologies for macaque (Reveley et al. 2017; Paxinos et al., 2024), marmoset (Paxinos et al. 2011), rodent (Swanson 2018; Wang et al. 2020) brains. Hierarchical levels were determined mainly based on brain regions, species and available gene markers. The harmonization process included regrouping of some structures such as OT and interstitial nucleus of the posterior limb of anterior commissure (IPAC), minor changes in structural acronyms and/or full structural description, as well as identification of presumed equivalent brain structures using consistent markers across species (**Fig. 3P**). For example, the OT in DHBA ontology was treated as a part of paleocortex (PalCx), which receives direct inputs from the olfactory bulb. In HOMBA ontology, OT is regrouped to STRv (**Fig. 3P**) based on most modern literature. Examples of minor changes include harmonizing acronyms (e.g., using all lower-case letters for all white matter structures), and using PVT and PVH for paraventricular nucleus of the thalamus and hypothalamus, respectively. For structures with two or more alternate names (e.g., fastigial nucleus of the cerebellum is also called medial nucleus in some literature), the second name is always placed in parentheses in the full name description in HOMBA [e.g., fastigial (medial) nucleus]. For orientation-related descriptions of brain regions and/or subdivisions we use rostral for most of the rostral, oral or anterior subdivisions in the literature, and caudal for most of the caudal and posterior subdivisions. Another orientation-related example is the harmonization of the dorsal and ventral lateral geniculate nuclei (DLG and VLG, respectively) across species. Although both DLG (or LGd) and VLG (LGv) are commonly used terms across species (particularly in rodents), other terms such as lateral geniculate nucleus (LG) and pregeniculate nucleus (PG) are also used in the literature, particularly in human and nonhuman primates (NHPs), in which the PG/VLG is located dorsal-rostral to LG/DLG. Thus, if DLG and VLG were used in human and NHPs, it would suggest that VLG would be located dorsal-rostral (rather than ventral) to the DLG/LG, causing inconsistent orientation description across species. To avoid this kind of confusion, we chose LG and PG as the harmonized terms for DLG/LGd and VLG/LGv, respectively (for detailed structural list of the HOMBA, see **Table S1).**

The combined analysis of multimodal data on the STR and consistent anatomical landmarks across species supported the rostral-caudal subdivision of STRd (Pu and Ca) and STRv into three major subdivisions (CaH, CaB and CaTd for Ca; PuR, PuM and PuC for Pu; NAC-OT, IPAC and PuCv+CaTv for STRv). CaH (head of Ca) and PuR (rostral Pu) are located rostral to the anterior commissure (ac) bundle and underneath them is the NAC-OT (typical STRv) (**Fig. 3A, B, C**). The PuM (middle Pu) is located between the level where the commissure appears and the level where the Pu starts to protrude ventrally to form PuCv [ventral subdivision of the caudal Pu (PuC); **Fig. 3D, E, F**]. Underneath the PuM is the IPAC, which is a small region that often shows NAC-like gene expression. The PuC starts at the level where the PuCv appears and extends caudally until the caudal end of the Pu. PuCv always displays similar gene expressions to NAC-OT (e.g., *PDYN* in **Fig. 3G, H, I**). The CaB (body of Ca) ends at the level where the PuC ends, whereas CaT (tail of Ca) is the remaining part of the Ca, extending from the dorsal to ventral direction and then rostroventral direction to reach lateral amygdala. Other evidence for striatal subdivisions across species is shown in **Figs. S3** and **S4**. For example, a clear dorsal-ventral difference (with gradient between) of the marmoset STR is found in sequential *WFS1*-ISH-stained sections **(Fig. S3**). Additionally, a clear rostral-caudal difference or rostral-middle-caudal gradient of the macaque Pu is seen in sequential CB-IHC-stained sections with strong, moderate and weak CB staining in the PuR, PuM and PuC, respectively although strong CB labeling is seen in all Ca subdivisions along the rostral-caudal axis (CaH, CaB and CaT) (**Fig. S4**).

Based on ventral-dorsal difference/gradient in gene expression (e.g., **Fig. S3**), the concept of STRv can be extended to include the IPAC, PuCv and CaTv (together called extended STRv in this study), in addition to typical STRv (i.e., NAC and OT) since all these components are located in the ventral aspect of the STR and consistently display NAC-like gene expressions and connectivity [e.g., **Figs. 3** and **S3**; see (Cho et al. 2013; Griggs et al. 2017; Ma et al., 2025) for connectivity].

### Harmonized 3D reconstruction of BG subdivisions across species

HOMBA ontologies and parcellations for BG structures, including STR, other BG components (GPe, VeP, GPi), and some closely related structures (STH, SN and VTA; **Fig. 4**), were 3D reconstructed in new MRI-based probabilistic templates or CCFs. These included the Human Connectome Project brain template [HCP template, (Glasser et al. 2016; Elam et al. 2021)], Mac25Rhesus template (https://balsa.wustl.edu/study/ggDBg), Mac25Cyno template (https://balsa.wustl.edu/study/ggDBg) and MarmosetRIKEN25 template (https://balsa.wustl.edu/study/ggDBg) (for details see **Methods** section).

Specifically, 2D histology-based delineations of BG subdivisions (e.g., **Figs. 3, S3, S4**) were mapped to matched MRI slices in 3D brain templates of each species and then revised and smoothed in coronal, sagittal and horizontal planes to generate 3D BG atlases for each species (see **Fig. 3J, K, L** and **Fig. S4** C’, E’ and H’ for this process). As shown in **Figure 4**, the overall structural organization is strikingly similar across species.

### Linking cellular and anatomical ontologies

Single cell and spatial genomics data generated in HMBA provide an additional evidence layer for defining anatomical boundaries, although it was not clear to what degree transcriptome-based cellular distributions would match gross anatomical boundaries. As described in companion studies, single nucleus Multiome analysis was used to create a detailed cell type classification (Johansen et al. 2025), with spatial transcriptomics data providing the spatial architecture of these cell types across the BG in all three species (Hewitt et al. 2025). These rich data offer an independent dataset for corroborating or revising histology and cytoarchitecture-supported parcellations, and for further iterative subdivision of parent structures based on spatial and molecular data. To enable co-visualization of spatial data with HOMBA parcellations, we performed nonlinear registration between the MRI coordinate space and the slab face image that was used for spatial sample dissection. When available for macaque and marmoset, the individual’s antemortem MRI was used as an intermediate registration step. Annotated HOMBA structures were then overlaid on the individual sections of spatial transcriptomic data (**Fig. 5A**), providing regional boundaries approximately in alignment with HOMBA annotations.

**Figure 5.**
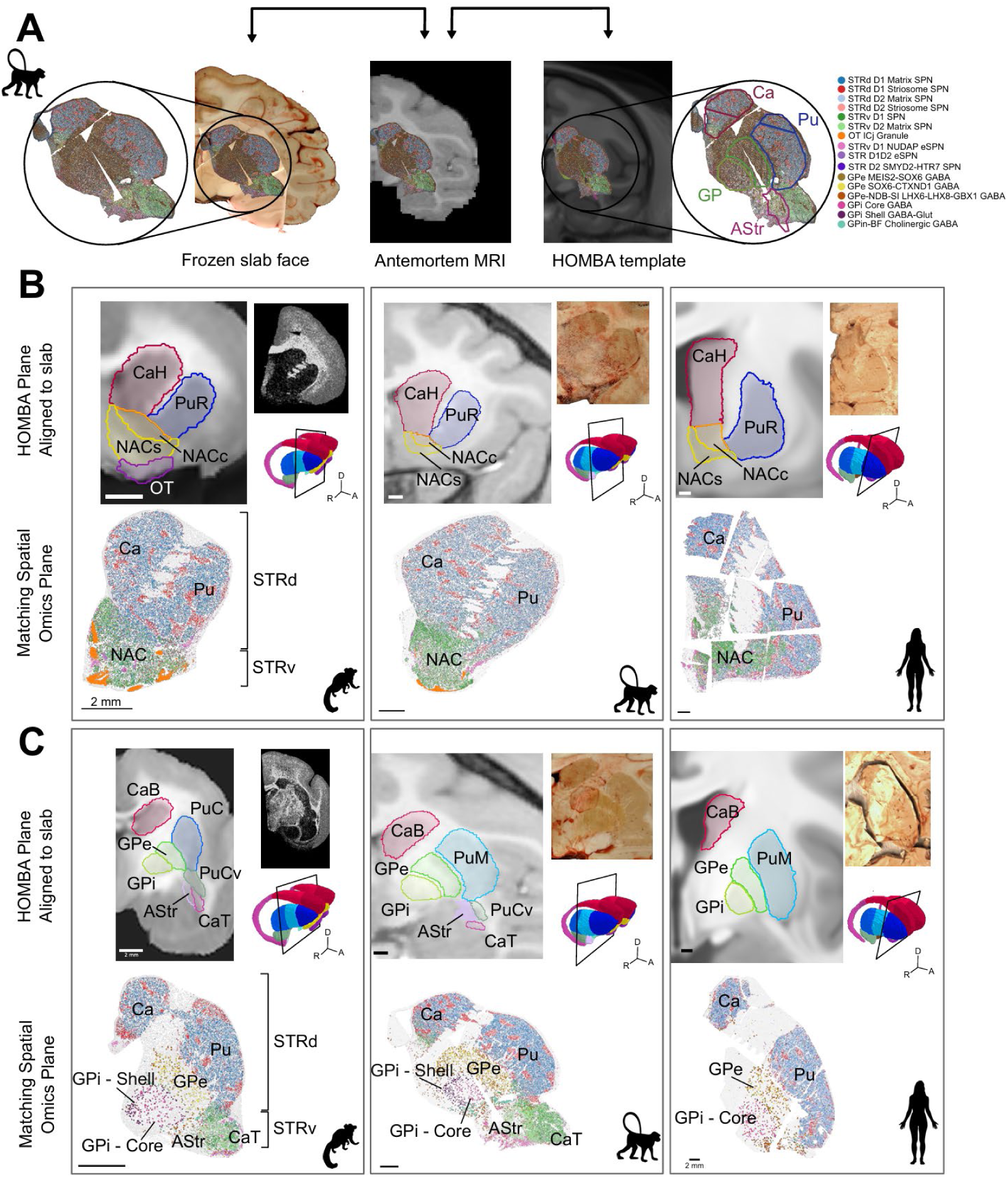
Spatial transcriptomic cell types and HOMBA parcellation boundaries. (**A**). Schematic diagram of registration protocol between HOMBA coordinate space and the dissected slab using the macaque as an exemplar. Serial transformations between slab face image, the individual’s antemortem MRI, and template MRI enable co-registration of HOMBA parcellations and spatial transcriptomic sections in the same space. (**B**). Comparison between HOMBA parcellations and cell type distribution in marmoset (left), macaque (middle), and human (right) rostral striatum. Top left, 2D planes aligned to slab face image with overlaid HOMBA annotations with 3D section context inset. Top right shows 2D slab face plane (macaque, human) or spatial transcriptomic section (marmoset). Bottom shows matched spatial transcriptomic sections with cells colored by cell **G**roup and HOMBA anatomical labels overlaid. (**C**) Comparison of HOMBA parcellations and spatial transcriptomic data in the caudal basal ganglia reveals finer anatomical divisions. HOMBA parcellation with MRI reference image registered to slab coordinate space (top) shows major anatomical divisions. Panels are organized in the same format as **B**. AStr, amygdalostriatal transition area, Ca, caudate nucleus, CaB, body of caudate nucleus, CaH, head of caudate nucleus, CaT, tail of caudate nucleus, GPe, external globus pallidus, GPi, internal globus pallidus, NAC, nucleus accumbens, NACc, nucleus accumbens core, NACs, nucleus accumbens shell, OT, olfactory tubercle, Pu, putamen, PuC, caudal putamen, PuCv, ventral caudal putamen, PuM middle putamen, PuR, rostral putamen, STRd, dorsal striatum, STRv, ventral striatum

Overall, spatial transcriptomics data showed cell type distributions that agreed with coarse anatomical divisions; for example, STRd and STRv were differentiable by their differential cell type composition (**Fig. 5B**), as were the GPe and GPi (**Fig. 5C**). However, there were also discrepancies. In the STRd, Ca and Pu had a similar distribution of cell types (primarily STRd D1 and D2 matrix SPNs) that did not show a regional boundary, suggesting that the Ca and Pu are anatomical divisions separated by the internal capsule but not differentiated by cellular makeup. Similarly in the STRv, the NAC had a uniform distribution of STRv D1 and D2 SPNs that did not reveal the core (NACc) and shell (NACs) subdivisions (**Fig. 5B**). Spatial cell type distributions at the confluence of the CaTv, PuCv, IPAC, and amygdalostriatial transition area (AStr) showed blurred boundaries that were not consistent with delineations of the HOMBA parcellations (**Fig. 5C**), but rather support grouping of these subregions into the extended STRv of HOMBA. In other cases, spatial transcriptomic data revealed subdivisions that were not identified using cytoarchitectural evidence alone. For example, GPi Core GABA cell types were spatially segregated from surrounding GPi Shell GABA-Glut cells (**Fig. 5C**), whereas these subregions of GPi, were only apparent in cell-type localization and are not currently represented in the HOMBA ontology.

There was obvious structurally restricted spatial organization of cell types across the basal ganglia despite the discrepancies noted above. **Figure 6** shows the qualitative distribution of cell types across structures in HOMBA BG structures, with cell type abundances rated as high, medium, and low levels across all three species, omitting cell types that either showed no presence in these areas or was too sparse to be confidently evaluated (below noise floor). This anatomical localization is a key part of the description of transcriptomically defined cell types, and was incorporated as part of the cell type nomenclature to denote anatomical localization, selective minimally necessary marker genes, and sometimes developmental origin or cardinal cellular properties (Johansen et al. 2025). This proved to be a challenge in matching two hierarchically organized ontologies, one structural and one cellular based on molecular similarities. On the cell type side, the mouse and human whole brain cell atlases (Siletti et al, 2023; Yao et al., 2023) have demonstrated a higher order hierarchical organization of cell neighborhoods, classes, subclasses and groups that often span a broad anatomical domain at higher levels and more specific structural restriction at lower levels. We used that knowledge of higher order organization in the cell type nomenclature even though it goes well outside of the basal ganglia structures. For example, the striatal GABAergic neurons are all in a broad neighborhood of Subpallial GABAergic neurons, with multiple classes restricted to the cerebral nuclei (CN) that are of known developmental origins from the medial (MGE), lateral (LGE) or caudal (CGE) ganglionic eminences. Another GABAergic class is distributed broadly across the Forebrain (F) and Midbrain (M), and is therefore called F M GABA. At the Group level, anatomical localization typically was more specific, restricted at the level of STR, STRd, GPe, etc. Most of the cell Groups in the striatum, comprising the CN MGE GABA, CN CGE GABA, and CN LGE GABA Subclasses, were broadly distributed across STRd and STRv and mostly did not localize to finer subdivisions of the HOMBA hierarchy (**Fig. 6**). Conversely, Groups of the F M GABA Subclass such as GPe SOX6-CTXND1 GABA and SN GATA3-PVALB GABA were specifically enriched in fine subnuclei such as GPe or SN respectively, and were named appropriately but also incorporating structure names outside of the basal ganglia if they were contained elsewhere (e.g. GPe-NDB-SI LHX6-LHx8-GBX1 GABA).

**Figure 6.**
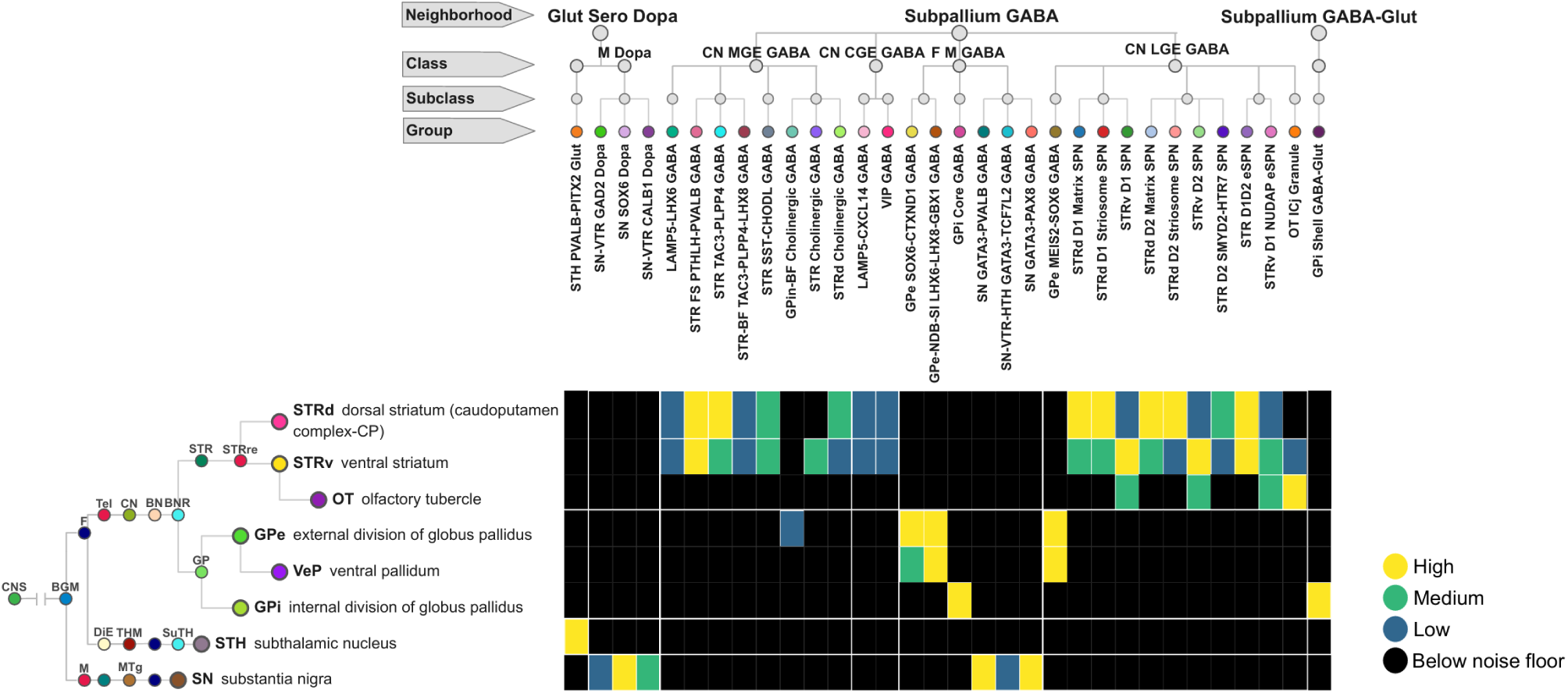
Cell types show complex spatial relationships across anatomical boundaries. Cell type by structure matrix showing qualitative scores of cell type **Group abundance** (horizontal axis) across HOMBA anatomical regions (vertical axis) in cross-species spatial transcriptomics datasets. Cell types that appear at levels below noise threshold are excluded. Light borders separate cell types by class and anatomical structures by major division. Note cell group names incorporate anatomical distributions.

### Gene expression topography supports parcellation of caudate subregions

Spiny projection neurons (SPNs), the main cell class of the caudate nucleus, showed cell group distributions that did not differentiate the HOMBA subdivisions CaH, CaB and CaT (**Fig. 6**). We therefore asked whether the human dataset showed systematic variation along the caudate at the finer level of granular cell type clusters and of gene expression within them. We analyzed pseudobulk expression profiles from sequencing libraries of 50 samples spanning the CaH (n=17), CaB (n=25), and CaT (n=8). Each library was assigned a 3D position in the HCP CCF (human) and projected onto a topographic axis capturing the rostrocaudal extent of the caudate (**Fig. 7A**). The relative abundance of SPN clusters varied substantially across subregions: 41 of 45 SPN clusters differed significantly in proportion across CaH, CaB, and CaT (Kruskal–Wallis, FDR < 0.05), and a random forest classifier trained on cluster proportions predicted anatomical region with 88% accuracy (**Fig. 7B; Fig. S5A**). The strongest spatial association was Human-30, a D1 Matrix cluster nearly absent in CaH (0.05%) but comprising 18.4% of SPNs in CaT (R² = 0.61).

**Figure 7.**
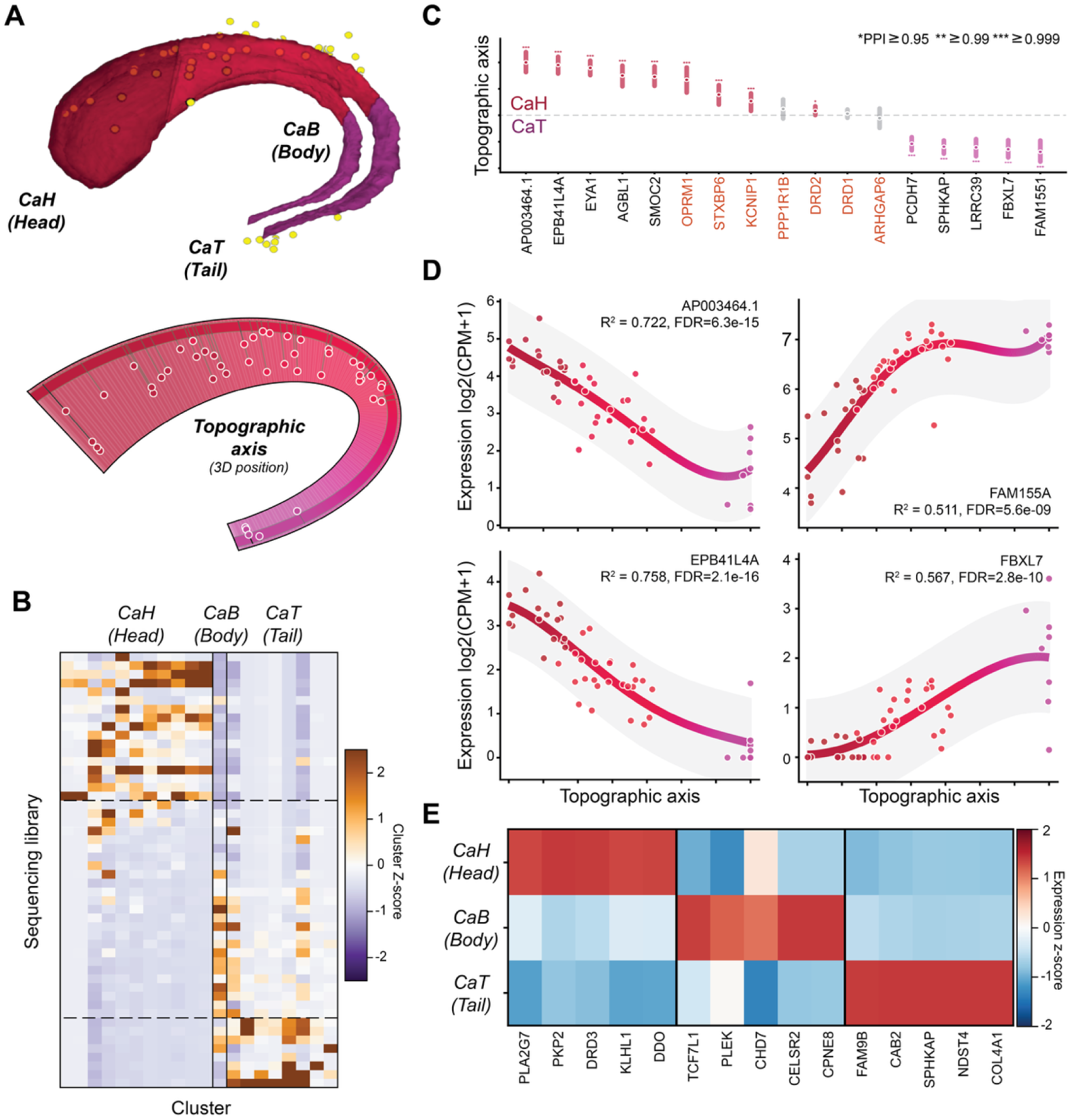
Gene expression and cell cluster variation in human caudate based on 3D coordinates of dissections. (**A**) Pinned sampling locations along the caudate longitudinal axis with CaH, CaB, and CaT boundaries overlaid. (**B)** Anatomical distribution of SPN clusters across the caudate subdivisions CaH, CaB, CaT. Cell clusters show anatomically restricted distributions. (**C**) Top variable genes in SPNs along the CaH-to-CaT axis. Many canonical SPN marker genes (*) show limited variation. (**D)** Expression versus topographic position, with line of fit from the models used to quantify variation; these are the data underlying the estimates in panel **C**. (**E)** Top marker genes for **SPNs** in three caudate anatomical regions.

We regressed pseudobulk expression against topographic position for each gene and SPN type independently (OLS, Benjamini–Hochberg correction). Across nine SPN types, 4,899 gene × cell-type combinations (2,483 unique genes) were significant at FDR < 0.05, ranging from 1,279 in STRv D2 SPN to zero in STRv D1 NUDAP SPN. In the pan-SPN pseudobulk data, 702 of 9,469 genes tested were spatially graded (FDR < 0.05). The strongest CaH-enriched genes, EPB41L4A (R² = 0.76), EYA1 (R² = 0.78), and AGBL1 (R² = 0.55), decreased monotonically from CaH to CaT, while the strongest CaT-enriched genes, FAM155A (R² = 0.51) and FBXL7 (R² = 0.57), increased along the same axis (**Fig. 7C, D**). Bayesian hierarchical modeling supported these effects: 35 of 46 genes modeled had 95% highest density intervals excluding zero (**Fig. 7C**). Notably, several canonical SPN markers showed no detectable spatial variation: DRD1 (FDR = 0.65), DRD2 (FDR = 0.22), PPP1R1B/DARPP-32 (FDR = 0.47), and ARHGAP6 (FDR = 0.76) were uniformly expressed along the axis (**Fig. 7C**, grey), consistent with the core transcriptional identity of D1 and D2 subtypes being maintained across the entire caudate even as hundreds of other genes vary with position. In contrast, DRD3 was strongly CaH-restricted (τ = 0.93, 14× enrichment; FDR = 4.0 × 10⁻⁴) and OPRM1 was significantly head-enriched (R² = 0.41, FDR = 5.7 × 10⁻⁵).

To identify markers for Ca subregions, we computed a specificity index (τ) for each gene across CaH, CaB, and CaT (**Fig. 7E**). CaH markers included PLA2G7 (τ = 0.97, 32× enrichment) and DRD3; CaT markers included FAM9B (τ = 0.85) and SPHKAP (τ = 0.82); CaB markers were more moderate, TCF7L1 (τ = 0.69), CHD7 (τ = 0.54), consistent with the caudate body representing a transitional zone. Gene ontology analysis indicated that CaT markers were enriched for heparan sulfate proteoglycan biosynthesis (adjusted p = 1.1 ×10⁻³; CSGALNACT1, NDST4, HS3ST1) and CaB markers for Wnt signaling (adjusted p = 2.9 × 10⁻³; TCF7L1, CELSR2, WLS).

Variance partitioning indicated that cell-type identity explained the majority of expression variance across all 444 pseudobulk samples, for 77% of genes, cell type contributed ΔR² > 0.05 beyond topographic position, while the reverse was true for only 5% (**Fig. S5B**, C). However, within individual cell type clusters the spatial signal was substantially stronger (**Fig. S5D**, E), indicating that the rostrocaudal gradient operates within each SPN type independently of inter-type differences. Comparison of individual CCF coordinate axes with the topographic axis showed that the CCF Y-axis alone was the best single predictor for 71% of genes, and the full XYZ model outperformed the topographic axis for 99% of genes tested (**Fig. S6**), suggesting that the three-dimensional CCF coordinates capture spatial expression variation at least as effectively as a derived one-dimensional projection. Overall, these results demonstrate both gene expression and cell cluster variation consistent with CaH, CaB and CaT parcellation, but also with more graded variation along the long axis of the caudate.

### Mapping and integration of multimodal BG data with macaque CCF

In neuroimaging, group average CCFs are routinely used as an anatomical substrate to aggregate and compare data across individuals. Data are accurately transferred, via volumetric and surface-based registration, from the individual to the CCF and from the CCF to the individual. Here, we have added another link in the registration chain for macaque, between an individual’s in vivo MRI space and ex vivo histological space. Using registration methods to be detailed elsewhere, the planar position of each tissue slab image within the T1w MRI volume and the in-plane in vivo vs ex vivo deformation was estimated via a semi-automated gradient-based optimization approach. This enables within-individual comparisons between functional and architectonic MRI and STx. In **Fig. 8A-C**, STRd and STRv cell types are colocalized with regions that are coactivated in different sensorimotor functional networks; a dorsal extremities network (top row) and ventral facial network (bottom row). This is a confirmatory finding, but it demonstrates the potential of the deeply integrated multimodal spatial framework. Furthermore, by concatenating the CCF-to-*in vivo* individual registration with an individual’s *in vivo* to *ex vivo* registration data can be precisely projected from the CCF to the individual’s histology and vice versa. For example, the underlaid structural annotations in **Fig. 8B** were drawn on the 3D CCF as described above, and projected to the individual macaque’s MRI space, then further projected onto each histological section. This permits the spatial distributions of cell types in individuals to quantitatively inform the placement of structure or substructure demarcations in the CCF for the iterative refinement of structural and cell type ontologies.

**Figure 8.**
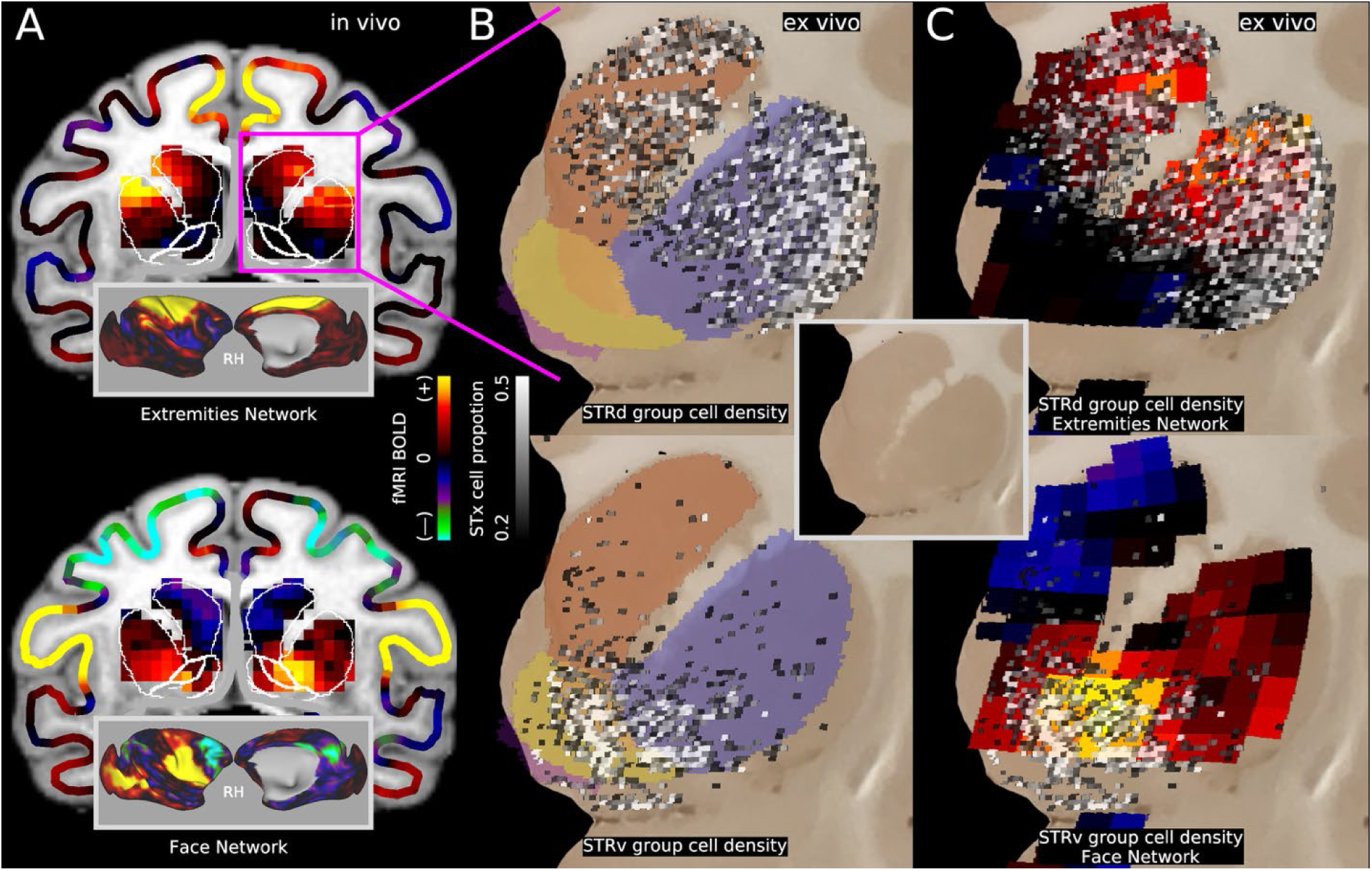
A joint spatial framework for MRI and STx within and across individuals. MRI and STx shown are from the same rhesus macaque exemplar individual with the MRI slice (A) precisely registered to the histological section (B,C). BG parcel annotations delineated on the Mac25Rhesus CCF are projected to individual *in vivo* MRI space as white outlines in A and individual histological ex vivo space, as colored underlay in B. (**A**) fMRI-derived temporal-ICA-based functional networks. Upper, Extremities sensorimotor functional network recruits STRd and limb neocortical effector fields. Lower, while facial sensorimotor functional network, recruits STRv and face neocortical effector fields. T1w structural images underlaid. Neocortical surface maps of networks inset. (**B)** STx cell type density. Upper, aggregate cell type proportion in 250μm bins of STRd D1 Matrix SPN, STRd D1 Striosome SPN, STRd D2 Matrix SPN, STRd D2 Striosome SPN cells. Lower, aggregate cell type proportion of STRv D1 SPN and STRv D2 SPN cells. (**C**) STx cell type density over functional networks projected into ex vivo space. Upper, STRd cell Groups and extremities functional network. Lower, STRv cell Groups and facial functional network. Section image without data overlay inset. Data at https://balsa.wustl.edu/M2l14.

**Figure 9.**
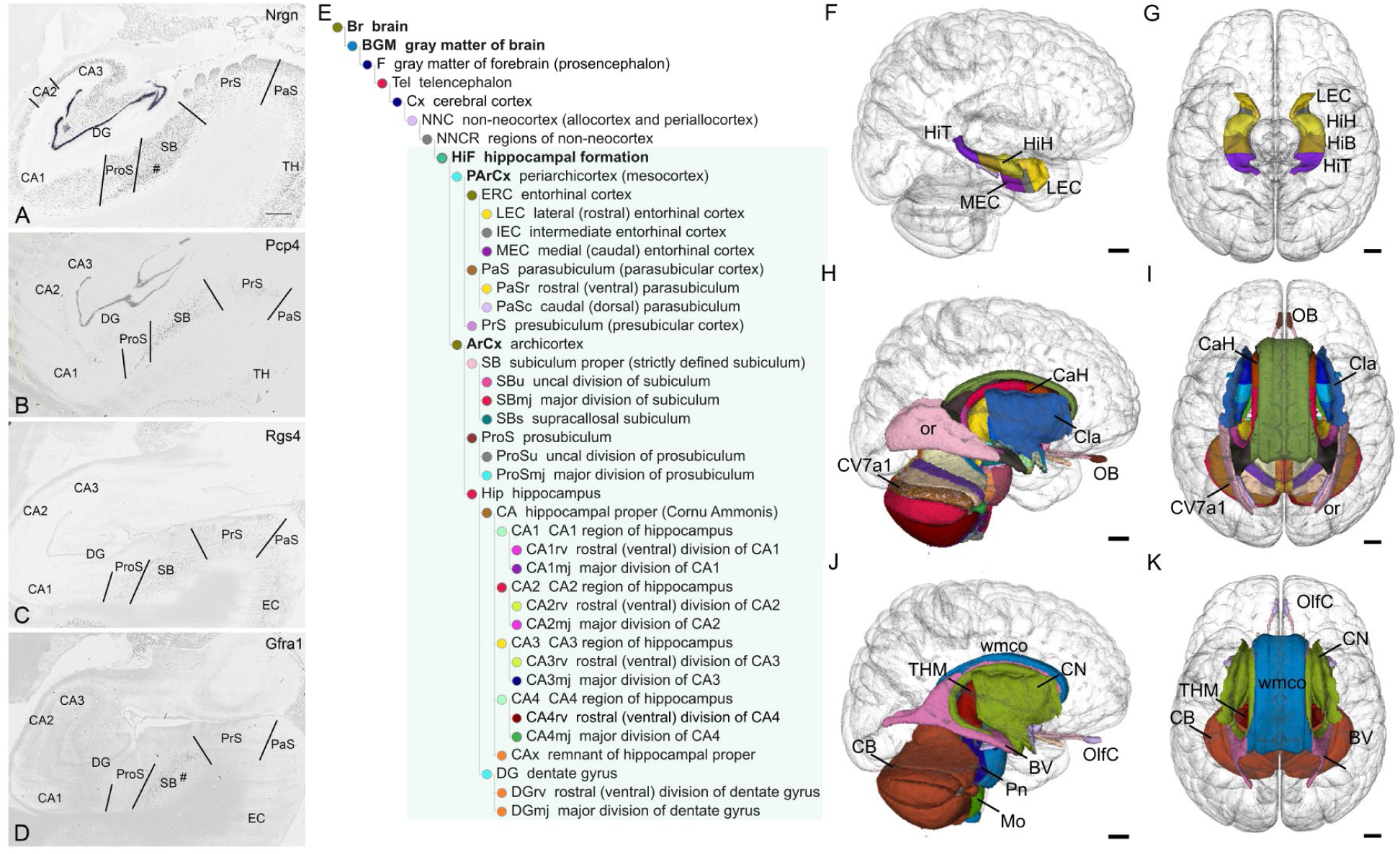
HOMBA annotation approach applied to human hippocampal and subcortical parcellations. (**A-D**) Marker genes *NRGN* (A), *PCP4* (B), *RGS4* (C) and *GFRA1* (D) used for annotation of the hippocampal subdivisions of the human ICBM2009b MNI template space. (**E**) HOMBA ontology highlighting hippocampal structural organization. (**F,G**) 3D parcellation of the hippocampal formation (HiF) in the human MNI template space with major divisions labeled. (**H-K**) Expansion of the annotation process to annotate 3D subcortical regions on the human template. (**H,I**) Sagittal and transverse views show the most granular structures supported by histological evidence. (**J,K**) Sagittal and transverse views show subcortical structures grouped by major divisions as defined in HOMBA. Error bars show 10 mm. BV: ventricles of the brain, CaH: head of caudate, Cla: claustrum, CB: cerebellum, CN: cerebral nuclei, CV7a1: lobule VIIaf, folium, HiB: body of hippocampus, HiH: head of hippocampus, HiT: tail of hippocampus, LEC: lateral entorhinal cortex, MEC: medial entorhinal cortex, Mo: myelencephalon, OB: olfactory bulb, or: optic radiation, Pn: pons, THM: thalamus, wmco: commissural white matter.

### 3D delineation and reconstruction of other brain regions using HOMBA

Using HOMBA and similar annotation approaches as for the BG, other brain regions can be parcellated and 3D-reconstructed consistently across species. For example, in the archicortex, which mainly include the hippocampus (DG and CA1-4), prosubiculum (ProS) and subiculum (SB), the boundaries of CA1, ProS and SB are often difficult to be consistently defined and delineated (Ding and Van Hoesen 2015; Ding, Yao, et al. 2020). Using HOMBA and a combined analysis of multiple markers, the borders between these three regions can be accurately and consistently delineated in each species even though the markers are not completely the same across species (**Fig. S7**). However, selection of multiple conserved region-specific markers is a key to accurate and consistent parcellation across species. In humans, gene *NRGN* is expressed in all three regions with strongest in SB while *PCP4* is mostly expressed in SB with weak or faint in ProS and CA1 (**Fig. S7A-B**). *RGS4* expression is mainly seen in ProS and SB with faint in CA1 whereas *GRFA1* is exclusively expressed in the SB with no/faint expression in both ProS and CA1 (**Fig. S7C-D**). When combining/aligning these four gene expression patterns, it is not difficult to consistently place the borders between the three regions. Similarly, in macaques (**Fig. S7E-H**), combination of the four gene markers (*NEFM, SYNPR, RGS4* and *GFRA1*) allows for consistent border delineation of CA1, ProS and SB. In marmosets (**Fig. S7I-L**, alignment of the expression patterns of the genes *CRYM, PCP4, RGD14 and GFRA1* also enables consistent border identification of CA1, ProS and SB. It is important to note the conserved *PCP4, RGS4* and *GFRA1* expression in CA1, ProS and SB across species. This combined approach also leads to consistent subdivisions of CA1, ProS and SB in mouse (Ding, Yao, et al. 2020) and of perirhinal cortex and temporal pole in humans (Ding et al. 2009; Ding and Van Hoesen 2010).

Using this cross-species analysis and the HOMBA ontology, the present study has delineated 280 subcortical gray and white matter regions and major ventricles in the 3D MNI152 human brain template (**Fig. G**H-K). Specific 3D-reconstructed structures are included in the annotation file: Developing Human Brain Atlas, version 2 (DHBAv2) — Common Coordinate Framework and Multi-Species Atlas Primer (CCF-MAP). Subcortical annotations and the HOMBA hierarchy can be used to group structures into 10 parent structures (olfactory cortex, cerebral nuclei, thalamus, midbrain, hypothalamus, cerebellum, pons, medulla, white matter, and ventricles) for aggregating spatial data (**Fig. G**J-K). Parent structures can also be used for masking and provide a resource for selecting subcortical structures of interest for downstream analyses (e.g., **Fig. S8**).

## DISCUSSION

We describe here a paradigm for the co-creation of human and non-human primate brain cell and brain structure atlases, and application of that paradigm in the BG. These atlases are intended to be foundational references for the community, creating a high-utility classification and map of cell types based on high information content single nucleus transcriptomics and epigenomics (snMultiomics) mapped to MRI-based spatial coordinate frameworks. This linkage bridges scales from molecular and cellular to structural and functional resolutions and harmonizes descriptions and analyses across species. Since the atlas is intended to describe and map distributions and tissue organization of cell types, its creation is necessarily iterative. Gross anatomy needs to be predefined to sample and analyze brain regions for snMultiomics to create the cell classification, and then refined with spatial transcriptomics for mapping into annotated CCFs. Creation of linked structural and cellular ontologies is an essential part of these cell atlases, as the structural distribution of cell types is a key element of cellular identity and function. In this way, mapping cellular distributions into MRI-based probabilistic templates or CCFs is a powerful means to analyze topographic variation and correlate fMRI signals to underlying cellular and molecular architecture, building on our prior human brain-wide gene expression atlas that has been a widely used resource by the MRI field (Arnatkevičiūtė et al. 2019; Richiardi et al. 2015; Burt et al. 2018; Hawrylycz et al. 2012).

The BG provides a relatively straightforward starting point, given its well defined structural architecture and species conservation across primates. Here, the HOMBA approach was successful in using existing literature, histological and gene expression information to delineate common structures across species, even when the same marker gene information was not available in all species. Spatial transcriptomics analyses based on much larger coordinated gene panels (Hewitt et al., 2025) generally substantiated the HOMBA structural parcellations as they became available. Structural nomenclatures canbe harmonized across species quite easily in the BG, with ontological mapping between HOMBA and other existing atlases, although generally HOMBA had finer subdivisions. For example, we subdivided the Pu and Ca into three parts with consistent anatomical landmarks such as anterior commissure (ac) and caudal edge of the Pu as reference points in combination with gradient gene markers and connectivity. The CaH, CaB and CaT of Ca are often reported as rough subdivisions of the Ca albeit no definition of the boundaries is described (Van Hoesen et al. 1981; Ding et al. 2016; Griggs et al. 2017). Interestingly, some NAC-like components, which always display similar gene expression to NAC, exist underneath the PuC (called PuCv), PuM (i.e., IPAC) and CaT (termed CaTv), are grouped as extended STRv, which was supported by spatial transcriptome data. In summary, the Ca, Pu and STRv can each be subdivided consistently into three basic parts (CaH, CaB and CaT for Ca; PuR, PuM and PuC for Pu; NAC-OT, IPAC and PuCv-CaTv for STRv), which are well in line with gene expression and connectional findings in NHP primates (see (Cho et al. 2013) for connectivity). Molecular and connectional findings from a recent study in mice also support these basic subdivisions of the STR (Ma et al. 2025).

Comparisons of cellular distributions to structural parcellations revealed a complex set of relationships. While the cellular organization supported major structural divisions like STRd versus STRv, other divisions did not correlate perfectly with cellular distributions. For example, the main spiny projection neurons (SPNs) located in the STRd did not cleanly vary between caudate and putamen. Rather, the division between caudate and putamen appears to be a structural division only, defined by the internal capsule that divides them. Many other aspects of cellular distribution and molecular variation are substructural, such as gradient and domain organization in Ca and Pu, striosomal organization in the Ca and Pu and islands of Calleja and OT clusters. Many of these features are highly variable from section to section and presumably individual to individual. Importantly, gradient organization appears to be a consistent substructural feature, but whether divisions such as the CaH, CaB and CaT reflect boundaries of divisions of more graded organization remains to be further investigated.

A major goal was to link cellular and structural ontologies and nomenclatures, as many cell types (and especially neuronal types) have anatomically restricted distributions. Where cell types had BG-restricted cellular distributions this was highly effective, and structural abbreviations were included in the cell type nomenclature. However, there are challenging nomenclature issues based on cell type distributions across the brain. The available whole brain single cell and spatial brain cell atlases and draft human brain cell atlas have revealed a higher order cellular organization based on molecular properties that very often span across structures. At one extreme, many non-neuronal cell types are relatively common across the entire brain; in contrast, some neuronal cell types are restricted to very specific brain regions. Linking structural and cellular ontologies therefore becomes an exercise in aligning two hierarchical ontologies built on different organizational principles, which requires understanding brain-wide hierarchical organization in both cases. There is a degree of symmetry between the hierarchical organizations of brain cell types and brain structures. Broader cell classes may be widely distributed (for example, across forebrain and midbrain), while more specific subclasses or groups within those classes may be quite anatomically restricted to specific structures. This alignment, and a principled and data-driven nomenclature system, will require iterative refinement as the whole atlas is created over time.

An important issue concerns the confidence with which homology assignments are made for structures in different species. Designating structure ‘A’ in the macaque as homologous to structure ‘B’ in the marmoset implies that they evolved from the same structure in a common ancestor. Since the brains of the common ancestor are generally unavailable, a rigorous proof of homology is generally infeasible, but a strong case may be made if sufficient comparative data are available (S.-L. Ding 2023; Xiang et al. 2023; S.-L. Ding 2024). It is expected that some of the homologies proposed in the initial HOMBA pairings/assignments will be challenged and that alternate homology assignments will be proposed in the future. The fundamentally hypothetical nature of proposed homologies applies to both cell type and structural cross-species ontologies. In both situations the evidence for homology is generally strongest at coarse levels of granularity (high in the hierarchy – see comment above) and becomes less clear when discriminating fine anatomical substructure or transcriptional clusters (i.e., low in the hierarchy). Transcriptomic cell type mapping across species provides a strong piece of evidence for homology across species, as transcriptomically-defined cell types can be quantitatively aligned across species based on shared molecular signatures (or integrated across species from their very definition (Johansen et al. 2025). The highly similar cellular makeup and 2D and 3D spatial organization across species argues for homology, although the finest molecular clusters cannot easily be aligned across species. Nevertheless, careful version control and handling of relevant data and homology adjustments will be important.

One of the major goals of the HMBA is to bridge microscale transcriptomics with mesoscale measures of brain structure and function, creating linkage between cellular and systems neuroscience that has not previously been available at this level of integration. Here, we demonstrate that such integration is both feasible and informative by aligning several data modalities within well-registered common and donor spaces. For example, we have established a tight spatial alignment between mesoscale MRI-derived structural and functional measures such as T1w/T2w myelin content and somatotopic sensorimotor BOLD networks and cellular precision spatial transcriptomics within the same donor and transferred that information into the common template. The HOMBA ontology of brain structures provides an annotated substrate by which to organize and navigate these multimodal data across species and scales. These combined cellular-systems approaches are anticipated to be critical for future work aimed at understanding brain evolution, health, and disease. For the BG subnuclei, these alignments were accomplished in 3D volumetric space; however, looking forward to investigations of cortical structures with sheet-like anatomy, multimodal alignments will also incorporate manifold 2D surface spaces. The extension of multiscale multimodal alignment and iterative co-creation of cellular and structural ontologies beyond the basal ganglia to other subcortical and cortical structures will be described in future works, further expanding the HMBA’s capacity to unify cellular and systems neuroscience across the brain.

The ultimate goal of HMBA is to build harmonized 3D whole human, macaque, marmoset and mouse brain structural and cell atlases, using HOMBA structures mapped into MRI-based CCFs and annotated and refined with transcriptomically-defined cellular distributions. These brain atlases will support many integrative functions including unified brain sampling, data integration, result mapping, interpretation and comparison across species, ages and labs, as well as comparisons between anatomy, cell types and function. Finally, it should also be pointed out that other brain parcellations from different species and modalities (e.g., connectional and functional parcellations) can also be mapped to the atlases/CCFs generated in this study by mapping the individual ontologies to HOMBA or vice versa, mapping HOMBA to other individual ontologies. All future changes will be versioned, documented and released in the CCF-MAP portal (Atlas Primer — Common Coordinate Framework and Multi-Species Atlas Primer (CCF-MAP)).

## RESOURCE AVAILABIITY

### Lead contact

Further information and requests for resources and reagents should be directed to and will be fulfilled by the lead contact, **Ed S. Lein**.

### Materials availability

This study did not generate new unique reagents.

### Data and code availability

All histological data including ISH and IHC data and existing brain atlases used in this project are all from previous publications, and many of them are publicly available. Accession numbers and links to the datasets are listed in the Key Resources Table.

Any additional information required to reanalyze the data reported in this paper is available from the lead contact upon request.

The HOMBA ontology can be accessed from the main documentation page [LINK], and is also available as owl ontology files. A version is available to explore on the Ontology Lookup Service (https://cellular-semantics.sanger.ac.uk/ols4/ontologies/homba).

## Acknowledgements

This publication was supported by and coordinated through the BRAIN Initiative Cell Atlas Network (BICAN) (https://braininitiative.nih.gov/research/tools-and-technologies-brain-cells-and-circuits/brain-initiative-cell-atlas-network). BICAN is funded by the National Institutes of Health’s Brain Research Through Advancing Innovative Neurotechnologies® (BRAIN) Initiative. This work was funded by the Allen Institute for Brain Science and by the National Institutes of Health (NIH) grant UM1MH130981 (PIs: ESL and HZ). The authors thank the founder of the Allen Institute, Paul G. Allen, for his vision, encouragement, and support. Additional BICAN (NIH) awards supported packaging and release of the atlases and CCF: U24MH130918 (PI: SM, LN, TM) and U24MH130919 (PIs: MHa, CT). Additional support was provided by the Institut national de la santé et de la recherche médicale (Inserm) IRP grant 2021-Cortical_Connectome, the European Research Council (ERC) Advanced Grant ERC-2023-Adv 101142153 PREDICTION, and the Agence Nationale de la Recherche (ANR) grant ANR-24-CE37-5022-CONNECTOME (PI: HK), as well as the Japan Agency for Medical Research and Development (AMED) grants JP18dm0307006 and JP23wm0625001, and the Japan Society for the Promotion of Science (JSPS) grant JP22H04926 (PI: TH).

## Author contributions

Design of the study: SLD, EL, LN, BR, DVE, HZ, MG, WF; Creation of HOMBA: SLD; Data analysis, curation and interpretation: SLD, AB, BL, BR, CT, DOS, FY, MHu, NJ, TH, NJ, SS, TI, WF, YHa, YHo; DVE, EL; Brain template generation: TH, TI, BR, AU, YHa, YHo, JV, MG, DVE, HK; 3D reconstruction: SLD, BF, JR, PL, AB; Web visualization: AB, TM, LN; Ontology management: AB, CT, DOS, UB, PR; Project management: LK, CT, HZ, EL; Grant procurement (PIs): EL, HZ, HK, TH; Manuscript writing-original draft: SLD, EL; Manuscript writing-review & editing: AB, BK, BL, BR, CT, LN, MHa, MHu, MS, NJ, RH, SS, SLD, TB, TM, XPL, YF, DVE, MG, EL; Figure preparations: SLD, AB, BF, SS, MHa, MHu, NJ, BR; Consultation: YH, ZL, YHa, WF, DH, BL, MHu, MHa, MS, FY, MS, YF, XPL, RH, BK, TB.

## DECLARATION OF GENERATIVE AI AND AI-ASSISTED TECHNOLOGIES

No AI-assisted technologies were used in the present study.

## METHODS

### EXPERIMENTAL MODEL AND STUDY PARTICIPANT DETAILS

#### Animals

All histological data from the human, macaque, marmoset, and mouse brains used in this study are from previous publications and publicly available (see **Key Resources Table**). No additional animals were used.

### METHODS DETAILS

#### Dataset collection and analysis

Harmonization of brain structural ontologies and regional parcellations requires sequential sections from whole bran hemispheres and/or large brain regions such as amygdala, thalamus, cerebellum, brainstem, hippocampus and cerebral cortex, as well as many stains from the same brain blocks or hemispheres. Based on these requirements, existing 2D whole brain histological atlases which contain sequential sections for multiple, or many markers are included (see **Key Resources Table**). For human brains, these include a 2D whole human brain reference atlas (S.-L. Ding et al. 2016), which contains whole brain sequential sections stained for Nissl substance and for parvalbumin (PV) and non-phosphorated neurofilament protein (NFP) immunohistochemistry (IHC with SMI32 antibody), and a large set of Allen *in situ* hybridization (ISH) data, which contains 250-400 marker genes for major brain regions such as neocortex, hippocampal formation (HiF), thalamus, hypothalamus, amygdala, cerebellum and BG (Hawrylycz et al. 2012). For macaque brains, we mainly refer to three published large datasets which contain many sets of sequential sections. The first is the Allen ISH datasets for macaque brains (containing 50-100 marker genes for whole brain and/or the major brain regions; see (Bakken et al. 2016). The second is the MacBrain histological dataset, which contains sequential whole brain sections of rhesus macaque (*Macaca mulatta*), stained for up to 19 sets of histological markers for each brain hemisphere (Selemon and Duque 2016). The IHC dataset from cynomolgus macaque brain sections (NeuN, SMI-32, PV, CB and Myelin) was obtained with a method described elsewhere (Lei et al. 2025) and aligned to the Mac25Cyno template for visualization with Neuroglancer. For common marmoset (*Callithrix jacchus*) brains, the marmoset brain atlas (Paxinos et al. 2011) which contains Nissl-, AChE-, PV-, NFP-, and CB-stained sequential sections, and Marmoset Gene Atlas (Shimogori et al. 2018; Kita et al. 2021), which contains many sets of ISH-stained sequential sections, are our main reference resources for identification of marmoset brain structures (see **Key Resources Table)**. Finally, 3D high-resolution whole brain MRI images from single brains (Edlow et al. 2016) which are registered to the MNI 152 template are also important reference data for 3D reconstructions of human brain regions/subregions.

All these data are analyzed with a combined approach. More specifically, multiple and/or multimodal datasets in sequential sections were aligned based on closely matched sectioning levels, as demonstrated in Figure 3 and Figure S5. This combined approach is essential for accurate and consistent identification and delineation of the boundaries of specific brain structures and for harmonization of structural ontology across species and ages.

#### Creation of a harmonized structural ontology across species

To generate harmonized 3D whole brain atlases across different mammalian species, a single structural ontology covers whole brain structures, and different developmental stages are needed. For this purpose, we have generated the HOMBA (online link), which contains 2,348 hierarchically organized neuroanatomical terms in the brain and spinal cord. These terms are mainly derived, modified and harmonized from the existing brain structural ontologies for human (S.-L. Ding et al. 2016; Mai et al. 2016; S.-L. Ding et al. 2022), macaque (Reveley et al. 2017; Paxinos et al. 2024), marmoset (Paxinos et al. 2011), and rodent (Swanson 2018; Wang et al. 2020; Paxinos and Franklin 2001) brains. The harmonizing process includes minor changes of structural acronyms and/or full structural description, combination of some structural features from different ontologies, as well as presumed identification of equivalent brain structures using conserved histological and/or molecular markers across species (see examples described in the Results section).

To generate an OWL ontology, we used a LinkML-OWL template to convert an Allen Brain Atlas structuregraph JSON representation* of HOMBA file to OWL, adding mappings to Uberon ported from an existing DHBA mapping to Uberon for all HOMBA terms with a direct DHBA equivalent. The ontology includes cross-references to DHBA and links to 3D reference images on the Allen Brain Atlas. Releases are available from https://github.com/brain-bican/harmonized_ontology_of_mammalian_brain_anatomy_ontology and browsable at https://cellular-semantics.sanger.ac.uk/ols4/ontologies/homba.

#### 3-D Annotation process of brain structures

Parcellation of subcortical structures was based on a combined analysis of intrinsic signals from the brain templates and matched high-resolution (7T) single-brain template (Edlow et al. 2019), as well as available supporting histological data including IHC and ISH data, which were manually matched to individual MRI slices based on main macro- and micro-anatomical features (e.g., **Fig. S2**). With ITK-SNAP multi-plane and 3D viewers, each brain structure was first delineated at regular intervals across coronal planes and then across sagittal and horizontal planes. This “weaved’ structure was filled in, refined and smoothed by illustration specialists. Once a critical number of 3D reconstructions was reached, a brain-wide merging of individual and local structure groups was performed. After merging, minor overlaps and gaps were resolved to attain complete seamlessness and any outstanding structure-related “negative spaces” appropriately defined. As additional structures were built, specialized macros for merging and splitting individual files were utilized to facilitate group delineations and adjustments at local and global levels, respectively. All these annotations were first performed on one hemisphere, and then these 3D reconstructions were mirrored to the other hemisphere to render a symmetrically complete 3D brain atlas/CCF. Detailed protocols for 3D delineation of individual and whole brain structures using ITK-SNAP are similar to those used for the 3D Allen Mouse Brain CCFv3 (Wang et al., 2020). A similar approach is being used to annotate macaque and marmoset brain structures in species-specific templates using ITK-SNAP.

#### Delineation of major white matter bundles and the ventricular system across species

Unlike the gray matter, many white matter tracts do not display specific gene markers. However, many major white matter bundles and all the ventricular subdivisions can be identified directly on a high-resolution MRI template (Edlow et al. 2019) and then parcellated on the low-resolution MNI152 templates since the former has been registered to latter templates (see the bottom row of Figure S2). In this study, 3D reconstructed major white matter bundles in the MNI152 template mainly include anterior commissure (ac), midline portion of corpus callosum (cc), olfactory tract (olt), fornix (fx) and its subdivisions, mammillothalamic tract (mtt), optic radiation (or), optic tract (ot), optic chiasm (ox), mammillotegmental tract (mtg), stria medullaris of thalamus (smt), posterior commissure (pc), cerebral peduncle (cpd), superior, middle and inferior cerebellar peduncles (scp, mcp and icp, respectively), transverse fiber of pons (tfp), medial lemniscus (ml), medial longitudinal fasciculus (mlf), decussations of scp (xscp) and pyramidal tract (xpy).

Identification of these fiber tracts on the T1w and/or T2w MRI template slices is relatively easier because most of them display differential contrast over adjoining gray matter structures. Similarly, the ventricular structures including the lateral (LV), third (3V), fourth (4V) ventricles, cerebral aqueduct (Aq) and central canal of the medulla (cec), as well as the choroid plexuses in 3V, 4V and LV, can be 3D-reconstructed directly in the template. These ventricular components overall display dark intrinsic signals on the T1w MRI template with high contrast to surrounding regions. Furthermore, we have subdivided the LV into rostral horn (LVr), body (LVb), atrium (LVx), caudal horn (LVc), and inferior horn (LVi) in the MNI152 template.

#### Web visualization of 3D CCF across species

Atlases/CCFs for each species were packaged into versioned Neuroglancer atlas assets using a data model derived from the AtOM schema (Kleven et al. 2023). In this model, an atlas is defined as a set consisting of an average template image, an anatomical parcellation (annotation set), an explicit coordinate space defining origin, voxel size, and orientation, and a controlled anatomical terminology with structure identifiers and names. For each species, the template image volume was converted into OME-Zarr, which stores large volumetric data as multidimensional arrays that can be accessed in parallel and streamed in a browser (Moore et al. 2023). Corresponding parcellation volumes were converted into Neuroglancer “precomputed” meshes for each structure and labeled with its HOMBA name and a high-contrast display color chosen to visually distinguish adjacent regions. Labeled parcellations were then combined at the parent level using HOMBA to create meshes for every level of the ontology. All atlas assets were deposited in a public Amazon S3 object storage and exposed through a Neuroglancer instance.

For the human brain atlas/CCF, we generated two atlas instances following this same procedure: one covering the whole-brain annotation set in MNI152 ICBM2009b space, and one covering a high-resolution BG annotation set in HCP MNI152NLin6ASym space.

#### 3D Macaque, Marmoset and Human MRI templates

The Mac25Rhesus, Mac25Cyno and MarmosetRIKEN25 templates were created using T1w and T2w images obtained from 25 individuals from each species with protocols and scanning parameters optimized using an HCP-style approach developed in the non-human primate neuroimaging & neuroanatomy project (NHP_NNP) (Hayashi et al. 2021). The data was collected with a 3T MRI scanner (MAGNETOM PRISMA, Siemens Healthineers, Erlangen, Germany) at RIKEN BDR, Kobe, Japan as well as at SBRI – Primage (Cermep, Lyon, France), in combination with high-sensitivity multi-array 24-channel (macaque) and 16-channel (marmoset) radiofrequency coils (Rogue Research, Montreal, Canada) (Autio et al. 2020). Structural images were acquired at isotropic resolutions of 0.50 mm in macaques and 0.35 mm in marmosets. Preprocessing included static magnetic field (B0) distortion correction using spin-echo field mapping, an automated brain extraction using bet4animal from FSL (Jenkinson et al. 2012), a signal inhomogeneity correction using the T1w multiplied by T2w method (Glasser and Essen 2011). The species-specific, resolution-scaled non-linear warping was performed using the fnirt from FSL using T1w images. All group-wise non-linear warping process involves a ’drift’—large-scale translation, scaling, or distortion as seen in the MNI templates (Glasser et al., 2016). The macaque and marmoset group average templates were corrected for this spatial bias by determining the dedrifting warp field and applying it to each individual’s data, then averaging the re-resampled images.

The human S1200_HCP3T1071 template was created using T1w and T2w images obtained from 1,071 individuals collected as part of the Young Adult Human Connectome Project and the procedures use for its creation have been previously described (Glasser et al. 2013). We have elected to keep the volume template in-register with the MNI152 6th-generation nonlinear asymmetric coordinate space (MNI152NLin6ASym) and therefore, this template inherits the drift present in that coordinate system. For the human template, the inaccuracies imposed by drift-caused distortions are outweighed by the explanatory power of conforming to widely adopted standards.

The volume templates are available from Atlas Primer — Common Coordinate Framework and Multi-Species Atlas Primer (CCF-MAP) and BALSA urls,. The pipeline to preprocess and standardize individual data to the template are publicly available at https://github.com/Washington-University/HCPpipelines. The animals, procedures, and results for the creation of the Macaque and marmoset MRI templates will be described elsewhere, as will the neocortical surface reconstructions, surface templates, and analyses of functional data collected in these cohorts.

#### Individual MRI

As with the templates, these images were preprocessed with an NHP optimized HCP-style approach. In addition, Macaque 2 underwent more than 40 hours of anesthetized resting-state and awake cognitive task MRI with MION contrast. These data were preprocessed and denoised with the HCP pipelines. Temporal ICA was performed on the entire concatenated denoised timeseries to yield model-free distributed brain networks. Figure 8 shows two selected exemplar networks. The detailed preprocessing procedures, as well as additional networks and functional analyses will be described elsewhere.

#### Individual to Template MRI Registration

As in the creation of the templates themselves the skull-stripped T1w MR images of the macaques 1 and 2 and marmoset 1 were registered to the Mac25Rhesus and MarmosetRIKEN25 templates, respectively using fnirt from FSL. The resulting non-linear warp field was inverted using invwarp from FSL. The CCF annotation volumes were resampled into each individual’s space with wb_command -volume-label-resample from Connectome Workbench (Glasser et al. 2013).

#### Histological Slab to Individual MRI Slice Registration in Macaque and Marmoset

Unfixed coronal slabs were cut and photographed on either their anterior or posterior face. The slabs were frozen and re-photographed. Fresh and frozen slab face images were cropped, oriented and scaled to a uniform physical scale. The cut-face tissue was segmented from via a lasso tool and background and non cut-face tissue removed. Fresh and frozen slab face images were registered to each other (https://elastix.dev) allowing either image to be used for registration validation.

The frozen slab face images were registered to the T1w MR volume though an iterative semi-automated gradient descent-based optimization method. Each slab face image was modeled as a transformed volume slice. The transformation consisted of a global affine component, which maps the volume slice into the plane of the slab image, followed by a nonlinear in-plane deformation. The affine component parameterizes slice position and orientation using translation, rotation, scaling, and shearing terms. The nonlinear component is represented by a dense 2D displacement field. To calculate this field, multiple displacement fields were applied at different spatial resolutions. Then, these sparse displacement grids were upsampled to image resolution, smoothed with a Gaussian kernel, and summed to produce the final dense deformation field. The transformation parameters were optimized to improve image similarity between each histological image and the transformed volume slice by maximizing the mutual-information between them. Regularization terms were employed to penalize deformation bending energy, mismatch between user-defined corresponding landmarks, and angular inconsistency between reconstructed slice planes, following an assumption of approximate coplanarity for adjacent slabs. The code for this 2D-3D registration algorithm, ‘deep-ssr’, is available from <github url>.

The registration of each slab face was manually evaluated by visually comparing resultant oblique T1w MRI plane and the in-plane outlines of the reconstructed white and pial surfaces the to the fresh or frozen slab face images in Connectome Workbench. For suspect registrations the automatically selected plane was corrected by manually adjusting the orientation of the oblique slice in a Multi-Planar Reconstruction rendering. The manually corrected planes were used to initialize subsequent algorithmic registration rounds. In some cases, the final automated registration step was limited to re-estimating the displacement field without re-optimizing the affine.

While the registration of the slab face images to the MRI slice was performed in the same way for each macaque and marmoset studied, additional registration procedures were dependent on the spatial transcriptomics format used for each individual. For macaque 1 (Fig. 5), slabs were cut into approximately 1 x 1 cm blocks before sectioning to accommodate the MERSCOPE format field of view. These block-wise sections were then imaged on the MERSCOPE platform and post-processed to derive cell coordinates and mapped cell taxonomies for each section. The resulting spatial sections were mosaicked as described in Hewitt et al. (2025). Briefly, spatial transcriptomic sections were registered to slab-face images with affine transformations using their corresponding block face images as a guide. This produces 2D coronal mosaicked spatial section planes in slab space. These 2D mosaicked planes were then registered to the MRI volume. Registered slab-face alignments provided initial estimates of rotation and anterior-posterior (A–P) position to guide this step. The position of each mosaicked spatial section plane was estimated as:

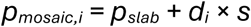

where *p_mosaic,i_* is the position of mosaicked spatial section plane *i*, *p_slab_* is the slab face position, *d_i_* is the depth of section *i* recorded from the cryostat during sectioning, and *s* is the scale factor to account for expansion from fresh tissue in the MRI imaging stage to frozen tissue in the blocking stage. The rotation of each mosaicked spatial section plane is shared with its slab face.

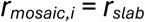

The parameters *p_slab_*, *r_slab_*, and *s* were jointly refined across all spatial section planes within a slab by comparing anatomical landmarks in each mosaicked spatial section plane with the corresponding MRI slice. A GUI visualization widget was created to visualize all mosaicked spatial section planes while adjusting the parameters *p_slab_*, *r_slab_*, and *s.*

In macaque 2 (**Fig. 8**), slabs were cut into approximately 2 x 3 cm blocks to accommodate the Stereo-Seq format and similarly mosaicked, though only one STx section was taken per block. For marmoset 1, the 1 x 2 cm Xenium format could accommodate the entire hemisphere for most sections and thus no mosaicking was performed (Hewitt et al. 2025). Registration of individual sections to MRI in marmoset 1 was performed using the same procedure as the slab face image to MRI registration described above, except that composite STx images of white and gray matter markers were used in place of the slab face photographs.

In all cases, the individual histology to individual MRI registrations were concatenated to the individual MRI to Template registrations to project individual histology into the CCF space. Inverted transformations were used to project 3D annotations from CCF space into the individuals’ MRI and histology spaces.

#### Histological Slab to Template MRI Slice Registration in Humans

Spatial transcriptomics sections were registered to their corresponding block-face images, and the block-face images were registered to slab tissue images as described in (Hewitt et al. 2025), producing mosaicked blocks comprised of continuous coronal planes. Each coronal plane was mapped to a corresponding 2D slice in the HCP 3T template associated with an anterior-posterior (A-P) index and slice angle (rotation along left-right and dorsal-ventral axes). The A-P index and slice angle was determined visually with landmarks on each coronal plane and its corresponding slab-face image (fresh and frozen), along with the recorded sectioning depth. Additional affine and nonlinear warp deformations between each coronal plane and its respective HCP 3T slice were estimated by performing ANTs SyN registration (metric=’CC’) between the manual anatomical annotations on each coronal plane and the corresponding HOMBA anatomical annotations. Initial affine registration parameters were estimated using Powell optimization-based affine estimation to maximize an average Dice coefficient across all masks. Masks that were small in area, such as the CaT and VeP were dilated and smoothed.

#### Spatial Transcriptomics

Spatial transcriptomics data was acquired from two macaque hemispheres, one marmoset hemisphere, and one human hemisphere. Macaque 1, marmoset, and human are included in a companion paper (Hewitt et al. 2025) with full methodological details. In brief, macaque 1 and human spatial transcriptomics data was generated using the Vizgen MERSCOPE platform across the full R-C extent of basal ganglia structures with a sampling interval of 1mm. Marmoset spatial transcriptomics data was generated using the 10X Xenium platform with a sampling interval of 200µm across the R-C extent. These experiments utilized 300 gene panels designed against the specific species and platform but with overlap where possible.

Macaque 2 spatial transcriptomics data was generated using the STOmics Stereo-seq platform. Unlike MERSCOPE or Xenium, Stereo-seq is a sequencing-based, full transcriptome assay. We utilized the large 2×3 cm chips which were able to tile a coronal hemispheric slab with 2-3 chips each. We sampled each slab once resulting in an R-C resolution of approximately 3-4mm. Data post-processing, including genomic and positional alignment was conducted using the Stereo-seq Analysis Workflow (SAW) (Gong et al. 2024). We then conducted a Cellpose-based cell segmentation workflow designed by the Allen Institute (SCALPEL) (Kunst et al. 2026) and was the same segmentation applied to macaque 1. Finally, basal ganglia cell type labels were transferred using MapMyCells (RRID:SCR_024672).

---

**Figure S1.**
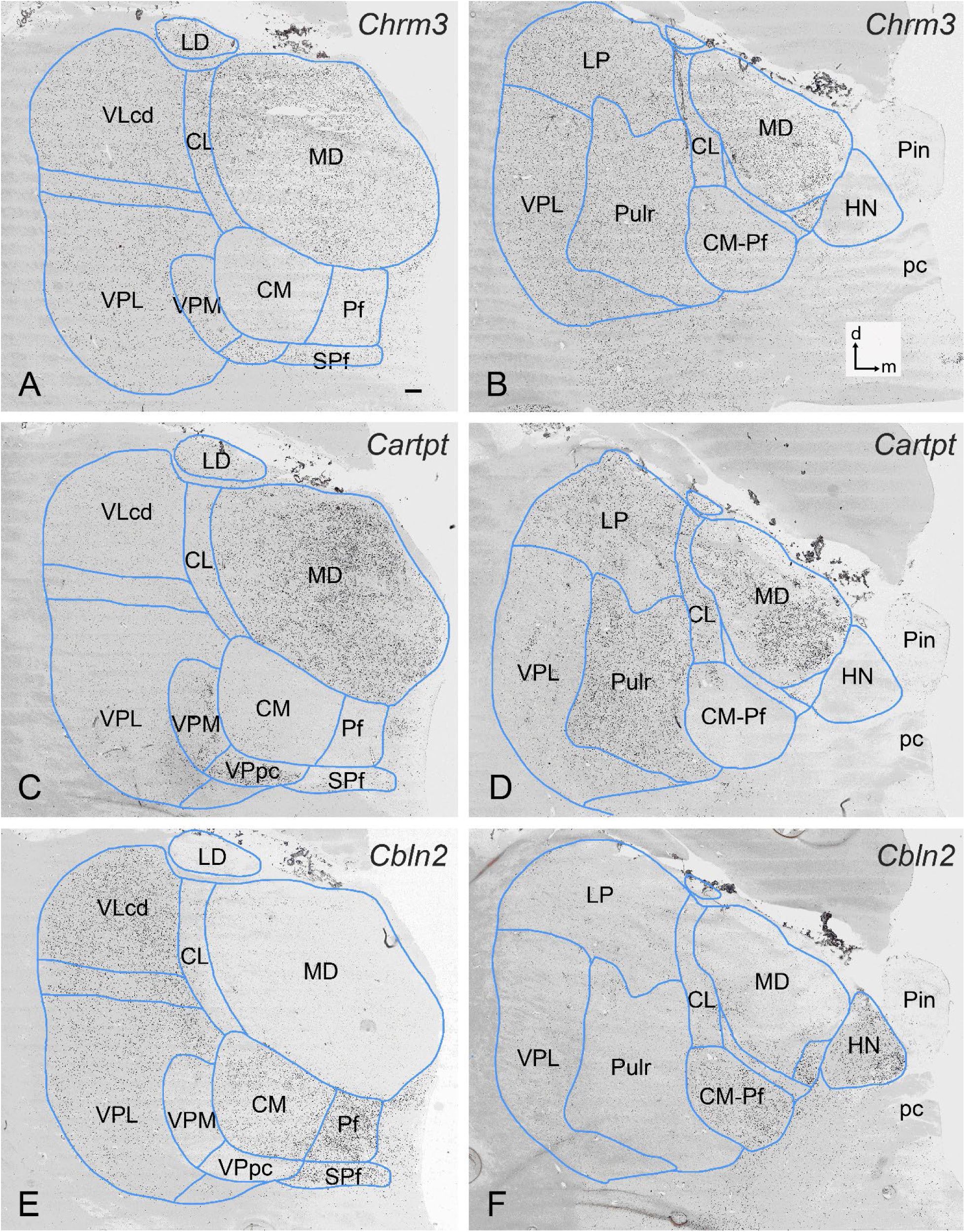
Delineation of human thalamic nuclei using a combination of multiple ISH-gene markers, related to Figure 2. (**A, B**) Cholinergic receptor, muscarinic 3 (*CHRM3*) expression in many human thalamic nuclei with weak/faint expression in the centromedian (CM) and habenular (HN) nuclei. (**C, D**) Strong CART prepropeptide (*CARTPT*) expression in the mediodorsal nucleus (MD), ventroposterior parvicellular nucleus (VPpc), latero-posterior nucleus (LP), and rostral pulvinar nucleus (Pulr) of the human thalamus. In contrast, ventral lateral nucleus (VL, VLcd), ventroposterior lateral nucleus (VPL), CM-parafascicular nuclei (CM-Pf), and HN have only weak/faint expressions. (**E, F**) Strong cerebellin 2 precursor protein (*CBLN2*) expression in the VL (VLcd), Pf, CM-Pf and HN of the thalamus with weak/faint expression in other adjoining nuclei. Note that consistent delineation of these nuclei requires comparisons across these and/or other gene markers. Panels (A, C, E) and (B, D, F) are three matched sections from two antero-posterior levels of the human thalamus. Scale bars: 790µm in (A) for all panels.

**Figure S2.**
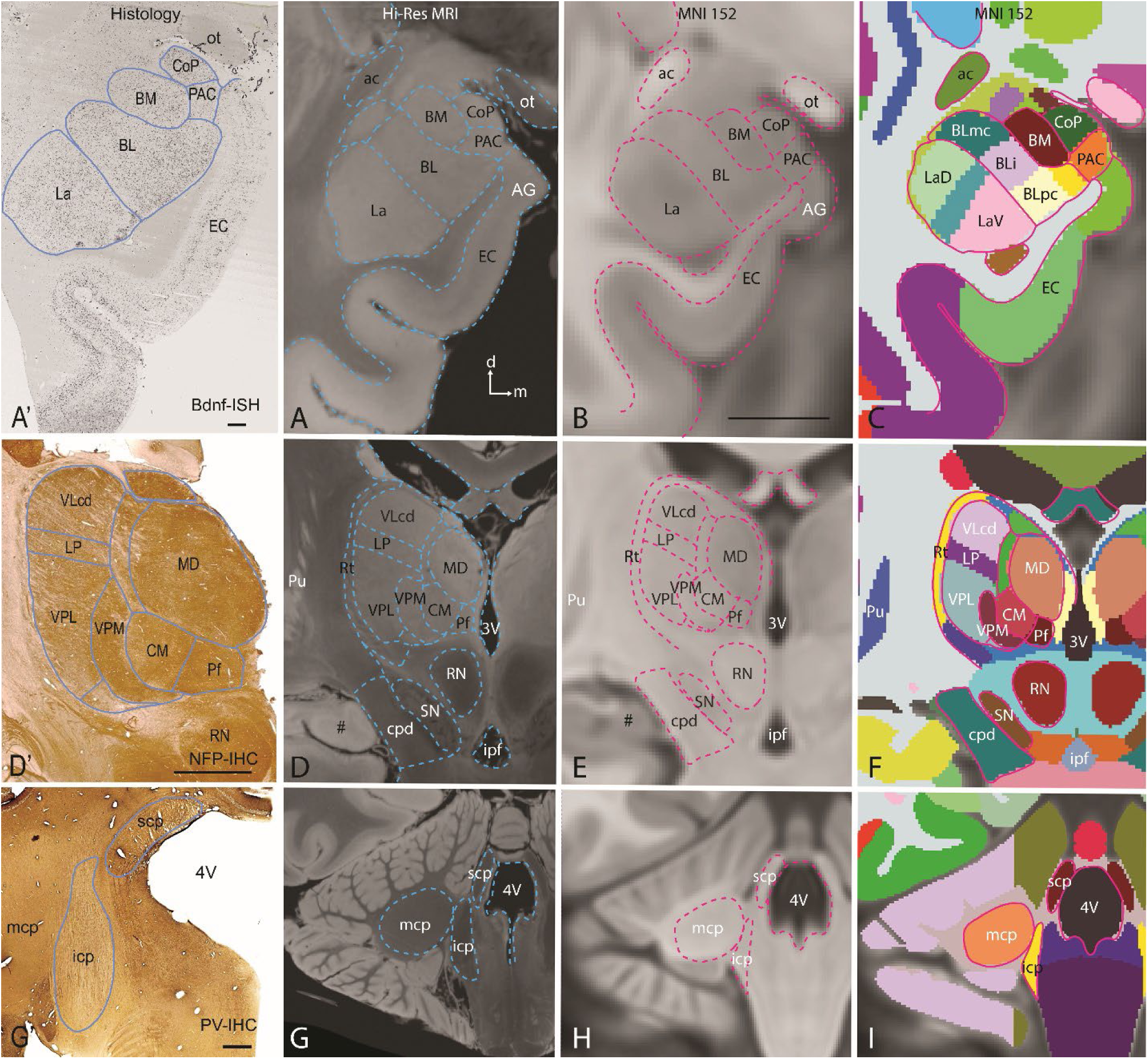
Mapping histology-based delineations to 3D human brain templates, related to Figure 2. (**A’, D’, G’**) Examples of matched histological sections to the MRI slices in panels (**A, D, G or B, E, H**), respectively. The histology includes ISH for the gene *brain derived neurotrophic factor (Bdnf <u>BDNF</u> in* **A’**), IHC for protein NFP (**D’**), and IHC for protein PV (**G’**). (**A-C**) Mapping histology-based delineations (**A’**) of the human amygdaloid nuclei to high-[**A**, T2w from (Edlow et al. 2019)] and low-(**B**, T1w from MNI152 template) resolution MRI slices by combining histological features and anatomical landmarks, which include anterior commissure (ac), optic tract (ot) and gyrus ambiens (GA). Some nuclei (e.g., BL) can be further subdivided into subdivisions (e.g., BLmc, BLi and BLpc in **C**) based on matched histological data. (**D-F**) Mapping histology-based delineations of the human thalamic nuclei using a similar approach. The landmarks used for matching the slices include the third ventricle (3V), interpeduncular fossa (ipf) and the most caudal putamen (Pu) and the intralimbic gyrus (indicated by #). (**G-I**) Mapping histology-based delineations of the human white matter tracts to the registered high-(G) or low-(H, I) resolution MRI slices from the 3D templates. The superior, middle and inferior cerebellar peduncles (scp, mcp and icp) are delineated based on anatomical locations or landmarks (e.g., G’) and a 2D human brain histological atlas (S.-L. Ding et al. 2016). Scale bars: 1600µm in (A’), 7080µm in (D’), 1550µm in (G’), 1cm in (B) for panels (A-I).

**Figure S3.**
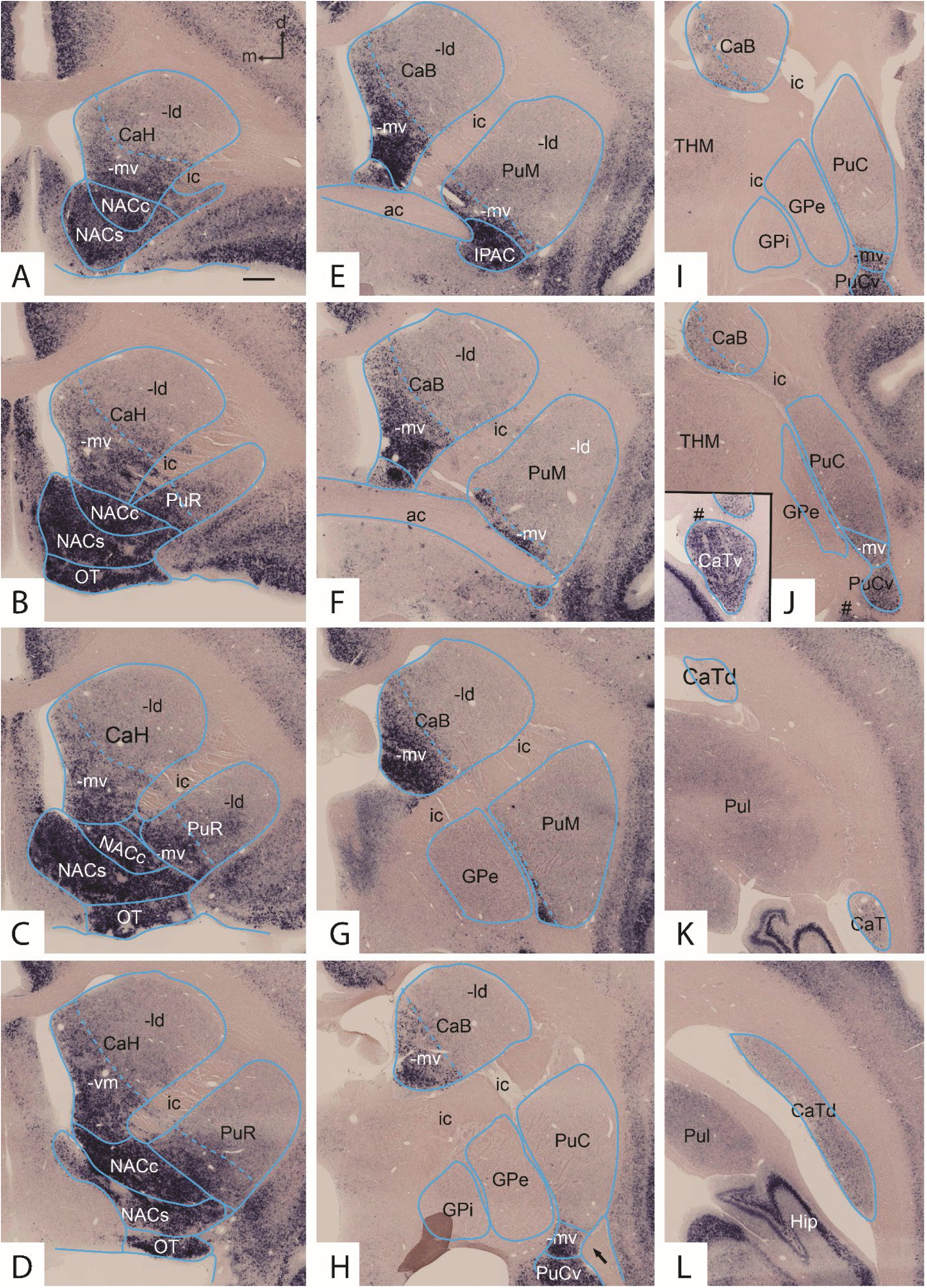
Example of dorsal-ventral striatal differences/subdivisions in the marmoset, related to Figures 2, 3, 4. (**A-L**) **Rostral-caudal** Wolfram syndrome 1 homolog (*WFS1)*-ISH-stained sections showing dorsal-ventral subdivisions and gradient organization of the striatum. It is noted that all the ventral striatal subdivisions along the rostral-caudal axis (rostral: NAC+OT; middle: IPAC; caudal: PuCv and CaTv) show strong *WFS1* expression. In addition, the ventral parts of the Ca (CaH and CaB) and Pu (PuR and PuM) show moderate (**A-C**) or strong (**D-G**) *WFS1* expression. At more caudal levels, the ventral part of Ca shows decreasing *WFS1* expression (**H-K**). Note that the dorsal and ventral parts of CaT (CATd and CATv, respectively) display weak and strong *WFS1* expression, respectively (see panels K, L and the inset in J). (**Inset in J**) Strong *WFS1* expression in the CaTv in the same section as the panel (J), in which the # indicates the corresponding point. Briefly, the *WFS1* expression pattern shows both ventral-dorsal and rostral-caudal differences in the striatum. It is worth mentioning that the expression pattern of the gene cannabinoid receptor 1 (*CNR1*) in the striatum (data not shown) is complementary to that of *WFS1* shown in this figure. Scale bar: 610µm in (**A**) for all panels.

**Figure S4.**
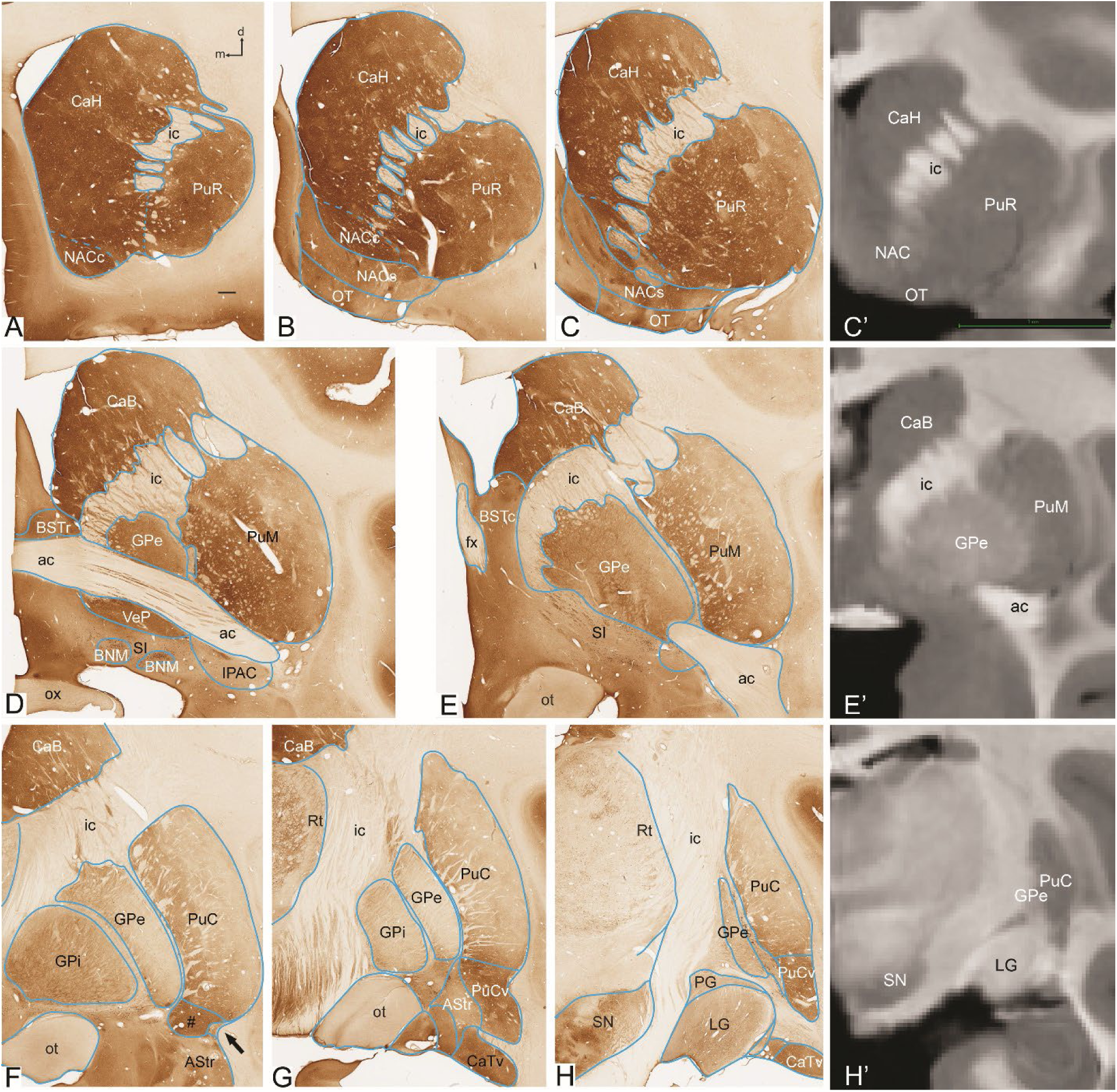
Example of rostral-caudal difference/subdivisions of macaque putamen, related to Figures 2, 3, 4. (**A-H**) Sequential calbindin-d28k (CB)-IHC stained sections show rostral (CaH-PuR)-caudal (CaT-PuC), medial (Ca)-lateral (Pu), and dorsal (Ca-Pu) and ventral (NAC-PuCv-CaTv) gradient or differences. To consistently subdivide the striatum (Ca, Pu and NAC-OT) along its rostral-caudal axis, the anterior commissure bundle (ac) is used as a landmark separating rostral and middle parts of the striatum. Specifically, the rostral Ca (i.e., head; CaH), rostral Pu (PuR) and rostral ventral striatum (NAC+OT) are located rostral to the commissure (**A-C**). The middle Ca (i.e., body; CaB), middle Pu (PuM) and middle ventral striatal part (IPAC) start at the level where the commissure appears (**D, E**). The caudal ending point of the CaB is at the level where the most caudal edge of the Pu is still clearly visible while the caudal point of the PuM is at the level where the lateral portion of the commissure located underneath the Pu (a little caudal to panel E) or before the medioventral part of the PuC (PuCmv, #) appears (F). Finally, PuC and PuCv start at the level where the lateral portion of the commissure is located lateral to the PuCmv (arrow in F) and extend caudally to the level where the caudal edge of the Pu appears. The CaT starts at the dorsal level where the PuC ends. CaT then travels ventrally and rostrally to reach the lateral amygdaloid nucleus (La) and lies ventrally to PuCv (**G, H**). Note the difference in the CB staining intensity between the PuR (**A-C**) and PuC (**F-H**) and between PuC and PuCv (**F-H**). (**C’, E’, H’**) Three MRI slices from the D99 macaque template that are matched to the CB-stained sections in panels (**C, E, H**) showing that the major BG subdivisions can be directly identified and parcellated in the MRI slices. Scale bars: 740µm in (**A**) for panels (A-H); 1cm in (**C’**) for panels (**C’**, **E’** and **H’**).

**Figure S5.**
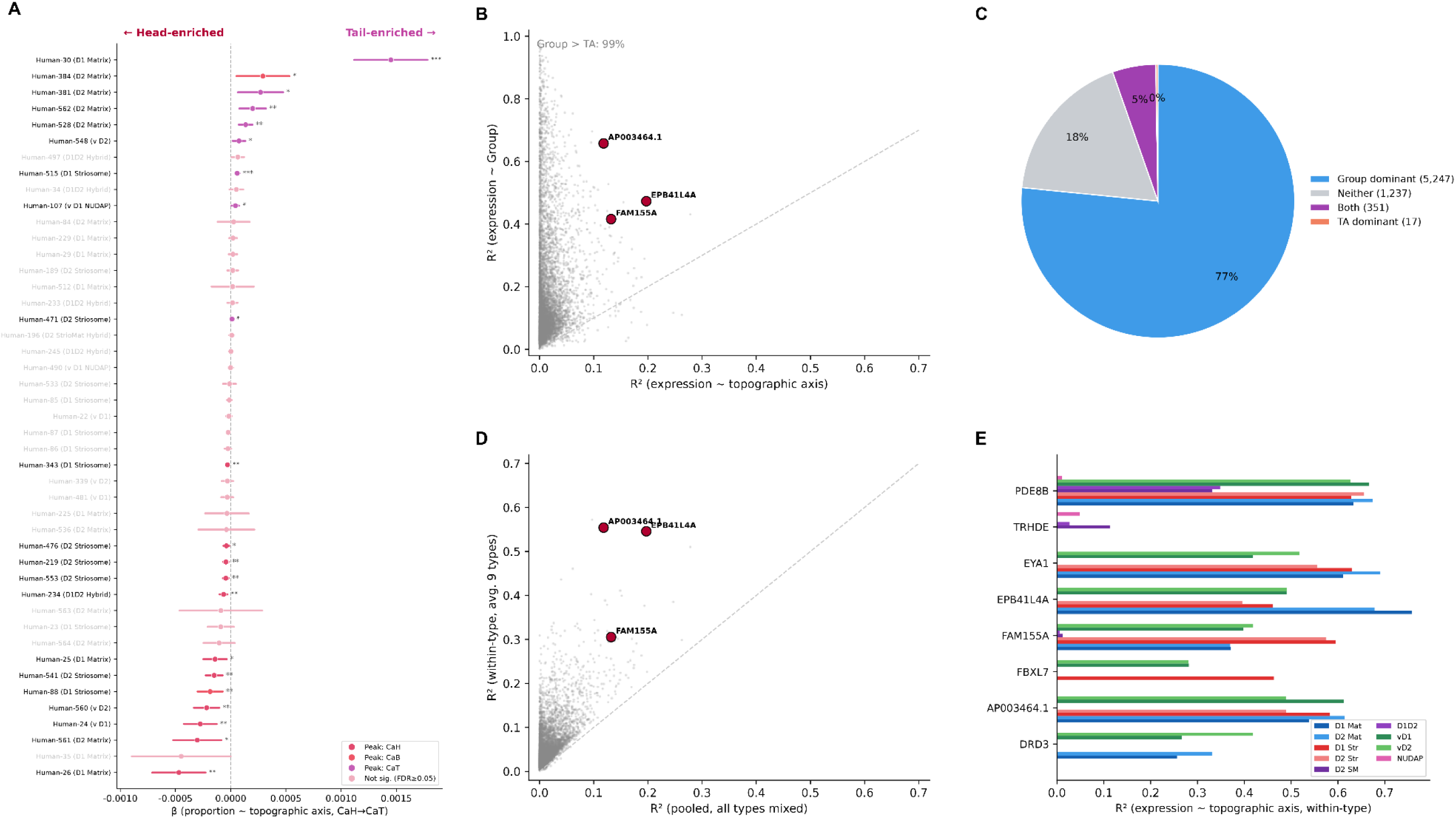
MSN cluster spatial proportions and variance partitioning, related to Figure 7. (**A**) Forest plot of the topographic-axis regression coefficient (beta, proportion ∼ topographic axis, CaH->CaT) for all 45 MSN clusters; points coloured by peak subregion, greyed where FDR >= 0.05; bars 95% CIs. (**B**) Per-gene R2(Group) vs R2(topographic axis) across all 444 pseudobulk samples (9 MSN types); Group exceeds topographic axis for 99% of genes. (**C**) Variance-partition category (Delta-R2 > 0.05): Group dominant, Neither, Both, TA dominant. (**D**) Within-type (averaged over 9 types) vs pooled topographic-axis R2 per gene. (**E**) Within-type topographic-axis R2 across the 9 MSN types for selected genes.

**Figure S6.**
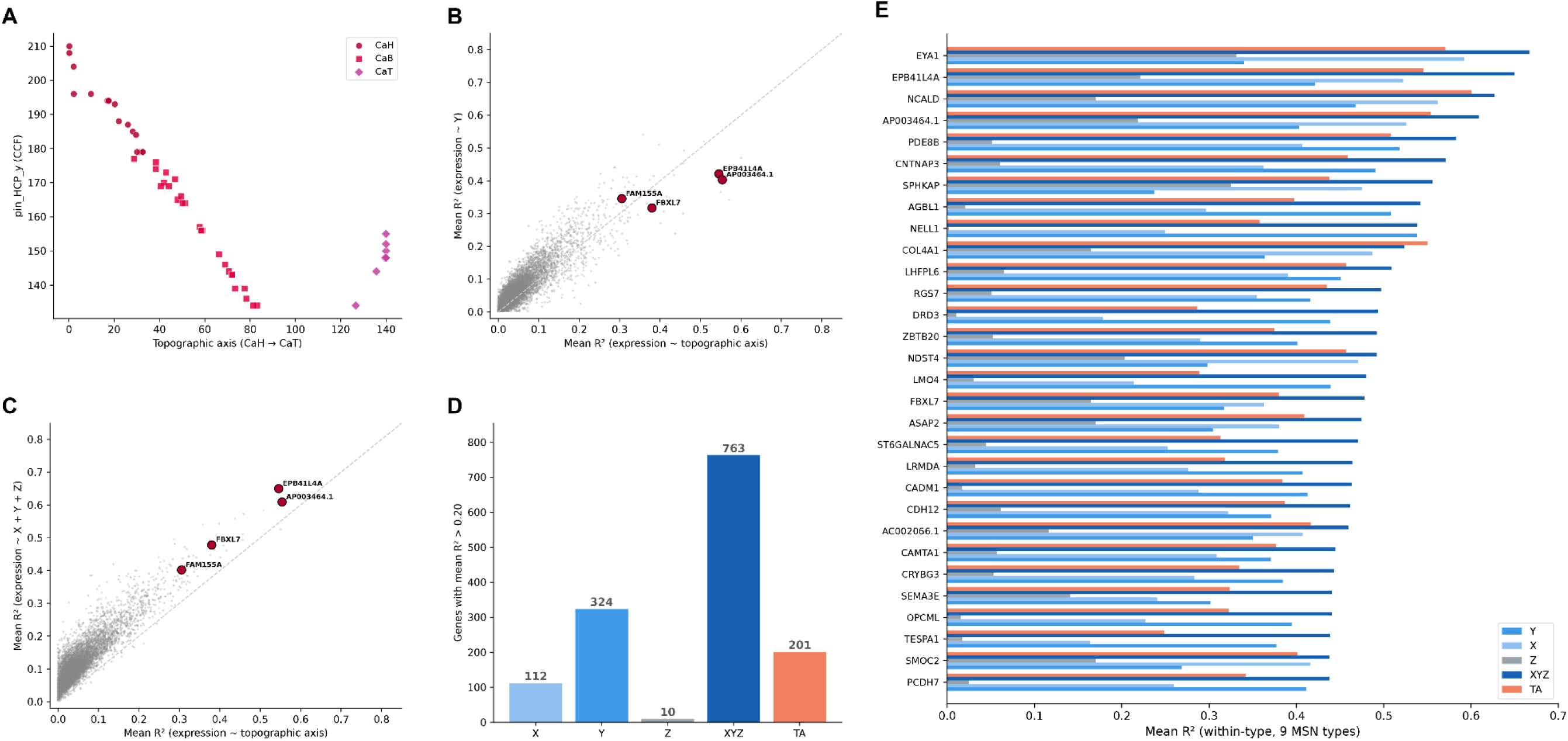
CCF-axis vs topographic-axis comparison and per-gene spatial variance, related to Figure 7. (**A**) Library pin_HCP_y (CCF) vs topographic position, colored by subregion. (**B**) Per-gene mean R2: CCF Y-axis vs topographic axis. (**C**) Per-gene mean R2: full XYZ model vs topographic axis. (**D**) Count of spatially detected genes per predictor (X, Y, Z, XYZ, TA) at mean R2 > 0.20. (**E**) Top 30 spatial genes ranked by XYZ R2, showing mean within-type R2 for each predictor.

**Figure S7.**
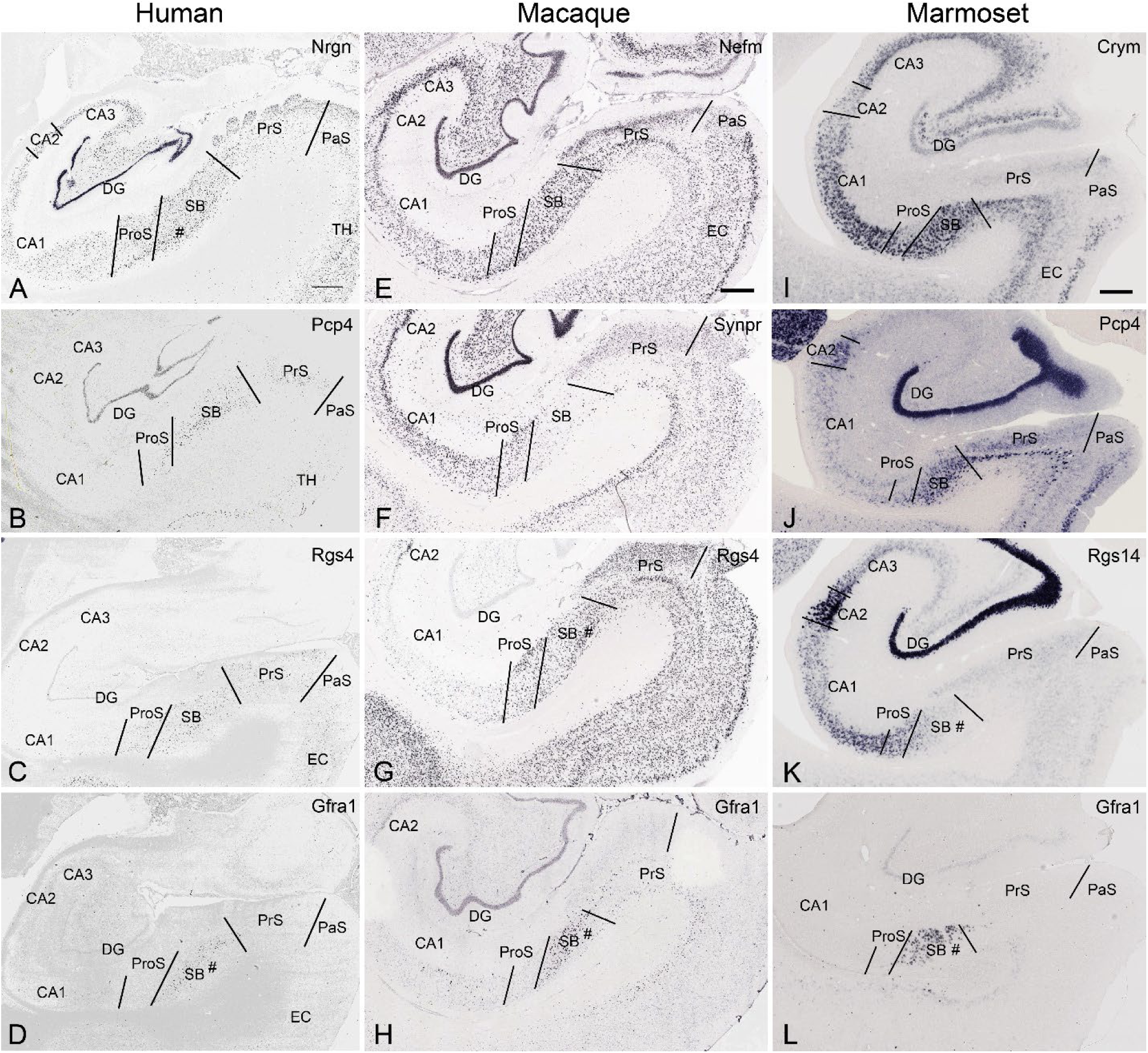
Histological comparison and harmonization of hippocampal subregions in human, macaque, and marmoset, related to Figure 8. (**A-D**) In human, expression of *NRGN* (A), *PCP4* (B), *RGS4* (C) and *GFRA1* (D) are used in combination for defining subregions and boundaries. (**E-H**) In macaque, expression of Nefm (E), Synpr (F), Rgs4 (G), and Gfra1 (H) are used. (**I-L**) In marmoset, Crym (I), Pcp4(J), Rgs14 (K), and Gfra1 (L) are used as markers for parcellation. Histological data is not available for some genes in specific species. Scale bars: 1600µm in (**A**) for panels (A-D); 800µm in (E) for panels (E-H) and 400µm in (I) for panels (I-L).

**Figure S8.**
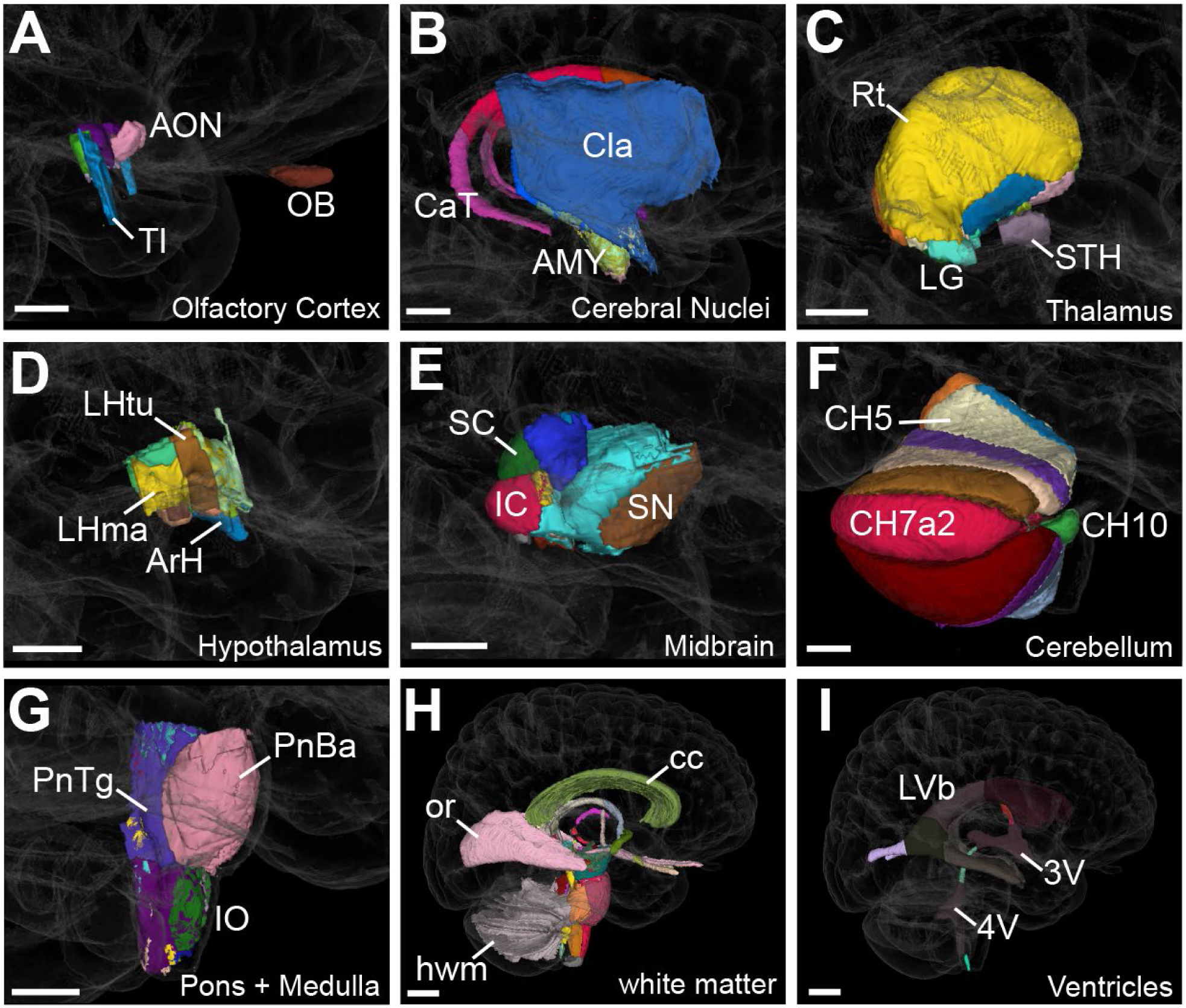
HOMBA parcellations of major human subcortical regions, related to Figure 8. The brain regions are indicated at the bottom right of each panel. Error bars are 10 mm. 3V: third ventricle, 4V: fourth ventricle, AMY: amygdaloid complex, AON: anterior olfactory nucleus, ArH: arcuate nucleus of hypothalamus, BV: ventricles of the brain, CaT: tail of caudate, cc: corpus callosum, Cla: claustrum, CH5: lobule V, CH10: flocculonodular lobe, flocculus part, CN: cerebral nuclei, CV7a1: lobule VIIaf, folium, hwm: hindbrain white matter, IC: inferior colliculus, IO: inferior olive, LG: lateral geniculate nucleus, LHma: lateral hypothalamic area, mammillary part, LHtu: lateral hypothalamic area, tuberal part, LVb: body of lateral ventricles, OB: olfactory bulb, OlfC: olfactory cortex, or: optic radiation, PnBa: basilar part of pons, PnTg: pontine tegmentum, Rt: reticular nucleus of thalamus, SC: superior colliculus, SN: substantia nigra, STH: subthalamic nucleus, TI: agranular temporal insular cortex.

